# Structural and temporal basis for agonism in the α4β2 nicotinic acetylcholine receptor

**DOI:** 10.1101/2022.02.23.481608

**Authors:** A. Sofia F. Oliveira, Isabel Bermudez, Timothy Gallagher, Susan Wonnacott, Giovanni Ciccotti, Richard B. Sessions, Adrian J. Mulholland

## Abstract

Despite decades of study, the structural mechanisms underpinning agonist efficacy in pentameric ligand-gated ion channels remain poorly understood. Here, a combination of extensive equilibrium and dynamical-nonequilibrium molecular dynamics simulations was used to obtain a detailed description of the structural and dynamic changes induced within the human α4β2 nicotinic acetylcholine receptor by a full and a partial agonist, namely acetylcholine and nicotine, and map how these rearrangements propagate within this receptor. These simulations reveal how the agonists modulate the patterns associated with intra and inter-domain communication and the evolution of the agonist-specific structural rearrangements. For the first time, we show that full and partial agonists, although generally using similar routes for through-receptor signal transmission, induce different amplitudes of conformational rearrangements in key functional motifs, thus impacting the rates of signal propagation within the protein. The largest agonist-induced conformational differences are located in the Cys loop, loops C and α1-β1 in the α4 subunit, loops F and β1-β2 in the β2 subunit and in the extracellular selectivity filter.

## Introduction

The α4β2 nicotinic acetylcholine receptors (nAChRs) are widely expressed in the brain where they influence cognition, mood, nociception and reward (*1–3*). The α4β2 receptors are members of the pentameric ligand-gated ion channel (pLGIC) family (*4*), which also includes the muscle nAChR, serotonin-type 3, glycine and γ-aminobutyric acid type A receptors. Mutations in the α4β2 nAChR can cause a rare familial epilepsy (*5*), and this subtype is necessary and sufficient for nicotine addiction (*1, 2*). The α4β2 subtype assembles in two distinct stoichiometries, namely the (α4)_3_(β2)_2_ or (α4)_2_(β2)_3_ (*6*). These receptors have a conserved pLGIC architecture consisting of a N-terminal extracellular domain (ECD), a transmembrane domain (TMD) formed by four transmembrane α-helices (TM1-TM4) that surround the ion-conducting pore, and an intracellular domain (ICD) connecting TM3 and TM4 (*4*) (Figure 1A). The agonist binding pocket is located at the extracellular subunit interface and is lined by six motifs: loops A-C from the (+) side and loops D-F from the (-) side of the interface (Figure 1B).

**Figure 1.**
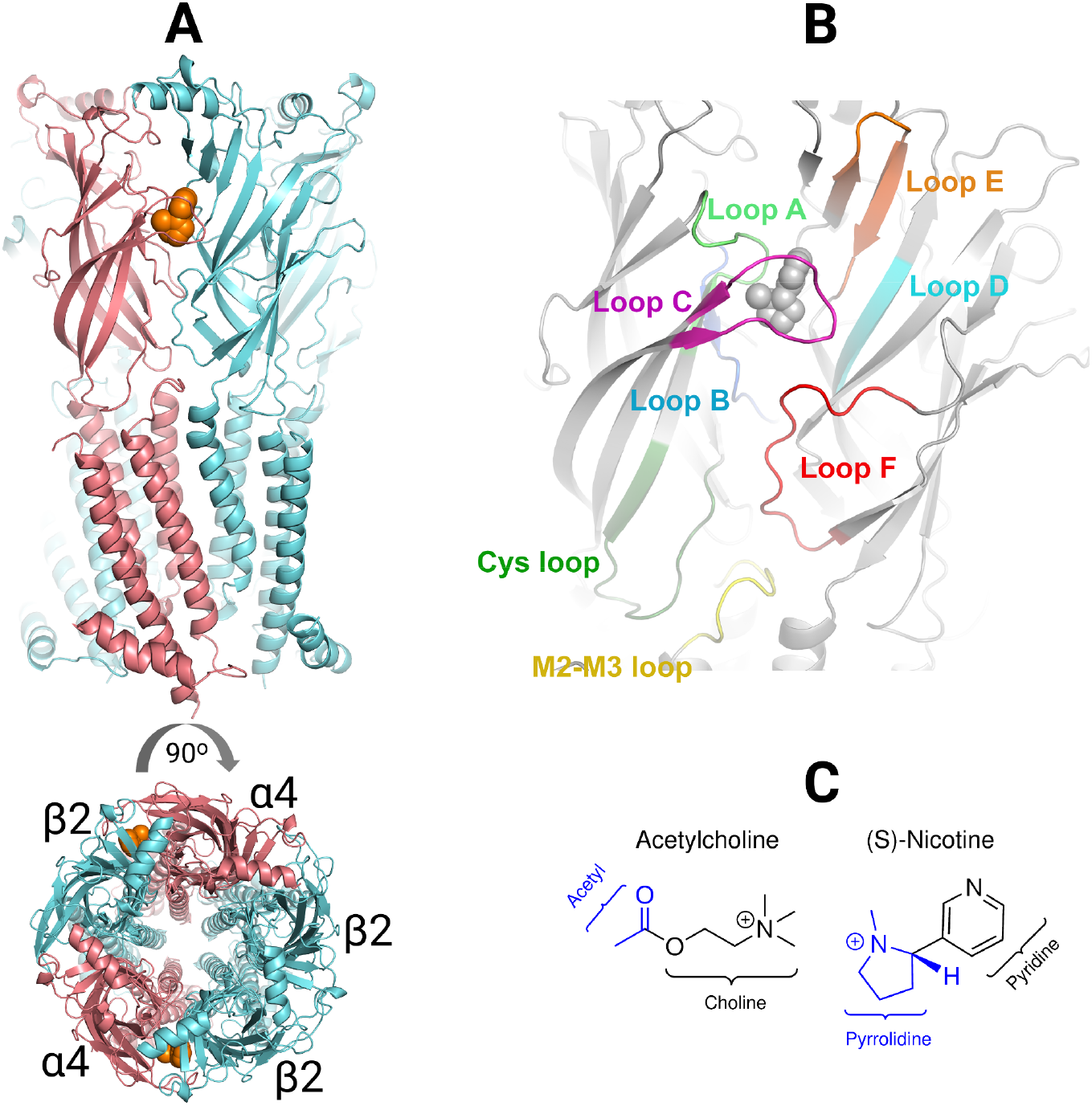
X-ray structure of the human (α4)_2_(β2)_3_ receptor. **(A)** Lateral and top view of the receptor (PDB code: 5KXI (*7*)). Note that this structure represents the (α4)_2_(β2)_3_ stoichiometry as the receptor is formed by two α4 (coloured in pink) and three β2 subunits (coloured in cyan). The orange spheres represent nicotine molecules occupying the two orthosteric binding sites. **(B)** Structural motifs involved in agonist binding and inter-domain communication. The motifs are highlighted with the following colour scheme: loop A, light green; loop B, blue; loop C, purple; loop D, cyan; loop E, orange; loop F, red; Cys loop, dark green; M2-M3 loop, yellow. Note that loops A, B and C from the (+) side and loops D, E and F from the (-) side form the agonist binding pocket. **(C)** Chemical structure of the simulated ligands: acetylcholine (full agonist) and nicotine (partial agonist). The different functional groups forming the ligands are highlighted with different colours.

Over the last decades, molecular dynamics simulations (e.g. (*8–15*)) and rate-equilibrium free energy relationships (REFER) (*16–19*) in combination with cryo-electron and X-ray structures of cationic and anionic pLGICs (e.g. (*7, 20–30*)) have shed light on the molecular interactions and structural changes that translate agonist binding to receptor activation. The mechanism for channel activation is broadly conserved across the pLGIC family and comprises an agonist-induced global twist in the ECD and TMD, leading to channel opening. The initial step in the activation process involves agonist binding to the orthosteric pocket(s) (*17, 18, 31*). It was shown experimentally that the binding of an agonists to a single orthosteric site has low probability of channel opening whereas occupancy of two non-consecutive sites enable proper activation of Cys-loop receptors (*32*). Agonist binding induces loop C closure (*33*) and a global conformational change that propagates to the ECD/TMD interface and then downwards to the transmembrane helices ultimately leading to gate opening (*12, 31, 34*). Activation of pLGICs is considered as a progressive stepwise isomerisation process proceeding from the agonist-binding pocket and to the TM2 helix (*12, 17, 18, 34*).

The molecules that activate pLGICs can be full or partial agonists, defined by the efficacy of maximally effective concentrations i.e. their ability to produce maximum responses (*35*). α4β2 partial agonists, in particular, are of therapeutic interest with some of them, namely varenicline and cytisine, being used as an effective strategy to aid smoking cessation (*36, 37*). Furthermore, there is compelling preclinical evidence that α4β2 partial agonists can also be useful for other brain disorders such as neuropathic pain and cognitive deficit (*38*).

Despite significant progress in understanding the activation of pLGICs, little is known about the structural basis of agonist efficacy in this family. Single-channel kinetic studies of different pLGICs, including nAChRs, performed at high temporal resolution have detected short-lived, preopen intermediates (termed flipped) states between the initial agonist-binding and the final pore-opening steps (e.g. (*39–42*)). It has even been proposed that partial agonism results from an agonist’s reduced ability to induce pre-gating conformational changes and achieve the flipped state, rather than an ability to partially gate the receptors (e.g. (*39, 41, 42*)). Recent structural studies of the glycine receptor suggest that partial and full agonists differ in their ability to induce the necessary conformational changes in the binding pocket, the event that initiates the transition of the receptors from the resting state to the flipped state and then to open (*43*).

Here, we have used a combination of equilibrium and dynamical-nonequilibriumcmolecular dynamics (D-NEMD) simulations to investigate the basis of partial agonism in the (α4)_2_(β2)_3_ subtype of the human α4β2 nAChRs and map the (structural and dynamic) differences induced by full and partial agonists in this receptor. In our previous studies, we have demonstrated how the combination of both equilibrium and nonequilibrium simulations is a powerful approach to study signal propagation and allostery in proteins (*44–49*), including nAChRs (*50, 51*). Now, a large set of D-NEMD simulations were performed together with extensive equilibrium simulations of the (α4)_2_(β2)_3_ subtype with either acetylcholine (full agonist) or nicotine (partial agonist) bound. The (α4)_2_(β2)_3_ stoichiometry was selected as it is the more prevalent, and displays high affinity for nicotine, which is well established as a partial agonist at α4β2 nAChRs (*52*), with less than 50% efficacy relative to acetylcholine (Figure S2). The approach used here not only allows for the mapping of the conformational rearrangements induced by each individual agonist but also clarifies how they impact signal propagation. We show for the first time that although full and partial agonists generally use a common pathway for signal transmission between the binding pockets and the ion pore, clear differences in the amplitude of the conformational rearrangements and in the rate of signal propagation exist between them. Such differences may be related to the efficacy of the agonists in this subtype.

## Results and Discussion

### Simulations of the acetylcholine- and nicotine-bound complexes

We performed extensive equilibrium MD simulations of the human (α4)_2_(β2)_3_ nAChR with two different agonists bound, namely the full agonist acetylcholine (ACh) and the partial agonist nicotine (NCT), to highlight the structural and dynamic differences induced by each in the receptor. Ten replicates, 280-ns each, were simulated for each agonist-receptor system in a total of 5.6 μs of simulation (see Supplementary Material for details). Each simulated system comprised the receptor inserted into an explicit lipid membrane (Figure S1), with two agonist molecules bound to the receptor (one in each binding pocket). Over the simulation time, all replicates remained stable with only less than 2% of secondary structure loss observed (Figure S4).

The dynamic behaviour of the agonists along the simulation time was monitored and, while ACh adopted many different binding modes inside the pockets (Figures S5-S6), NCT exhibited limited dynamics and generally remained in the same binding orientation (within both binding pockets) throughout the simulation (Figures S5 and S7). However, despite the differences in agonist mobility, the canonical interactions in both binding sites between the agonists and TyrA (Y100 in the α4 subunit), TrpB (W156 in the α4 subunit), TyrC1 (Y197 in the α4 subunit) and TyrC2 (Y204 in the α4 subunit) (*53–55*) are mostly present (Figures S8-S9). It is also interesting to note the differences in the distance distribution profiles between the agonists and the aromatic residues lining the pockets, mainly with TrpD (Figure S10). Intriguingly, and despite NCT being bulkier than ACh, the interaction formed with TrpB is on average shorter for the partial agonist. It was shown experimentally that common α4β2 agonists form a cation-π interaction with TrpB, which provides an anchor point for the ligands inside the aromatic box (*53*). The charged (i.e. protonated) secondary amine of NCT, in addition to this cation–π interaction, also forms a hydrogen bond with the backbone carbonyl of TrpB (*53*). The shorter TrpB-NCT distance observed in our simulations is likely due to this additional hydrogen bond, which is present 94% of the simulation time.

The analysis of the ACh and NCT dihedral angles shows that, despite their varying degrees of conformational flexibility, both ligands have preferred binding conformations inside the binding pockets (Figure S11-S12). The preferred mode for ACh corresponds to an extended arrangement (see Figure S11D for a representative example) with N-C-C-O dihedral≈65°, C-C-O-C≈175° and C-O-C-C≈175° (Figure S11). The most populated conformation of NCT has a dihedral angle between the pyridine and pyrrolidine rings of ≈ –65° (Figure S12), which is similar to the binding mode observed in the X-ray structure of the (α4)_2_(β2)_3_ nAChR-NCT complex (*7*).

### Agonist-induced structural and dynamic changes

The C_*α*_ root mean square fluctuations (RMSF) profiles were determined for the ACh- and NCT-bound systems to identify and characterise the dynamic changes induced in the receptor by the agonists. The average RMSF profiles for the two systems are very similar (Figures S13-S14), and no significant difference is observed for most of the structural motifs forming the binding pockets (namely loops A, B, C, D and E) nor at the ECD/TMD interface. Loop F is the only exception as it shows an increase in flexibility in the NCT-bound system. This motif is located at the bottom of the binding site and is a region with low sequence conservation among nAChRs. Voltage-clamp fluorometry experiments have shown that the upper part of the F loop is essential for ligand binding affinity and specificity whereas the lower part is thought to be part of the gating machinery (*56, 57*). Additionally, our previou work has also highlighted a complex contribution of this region to signal propagation from the binding pockets to the TMDs (*50, 51*). However, so far, the exact role played by the F loop remains unclear.

To identify the structural differences induced by full and partial agonists in the receptor, the Cα positional deviations between the ACh and NCT-bound systems were calculated as a function of the residue number (Figure S15). The final deviation values correspond to the average obtained over all the 100 combinations (resulting from the 10 ACh × 10 NCT pairs of trajectories). Those deviations were, afterwards, mapped on the average ACh structures (Figures 2 and S15).

**Figure 2.**
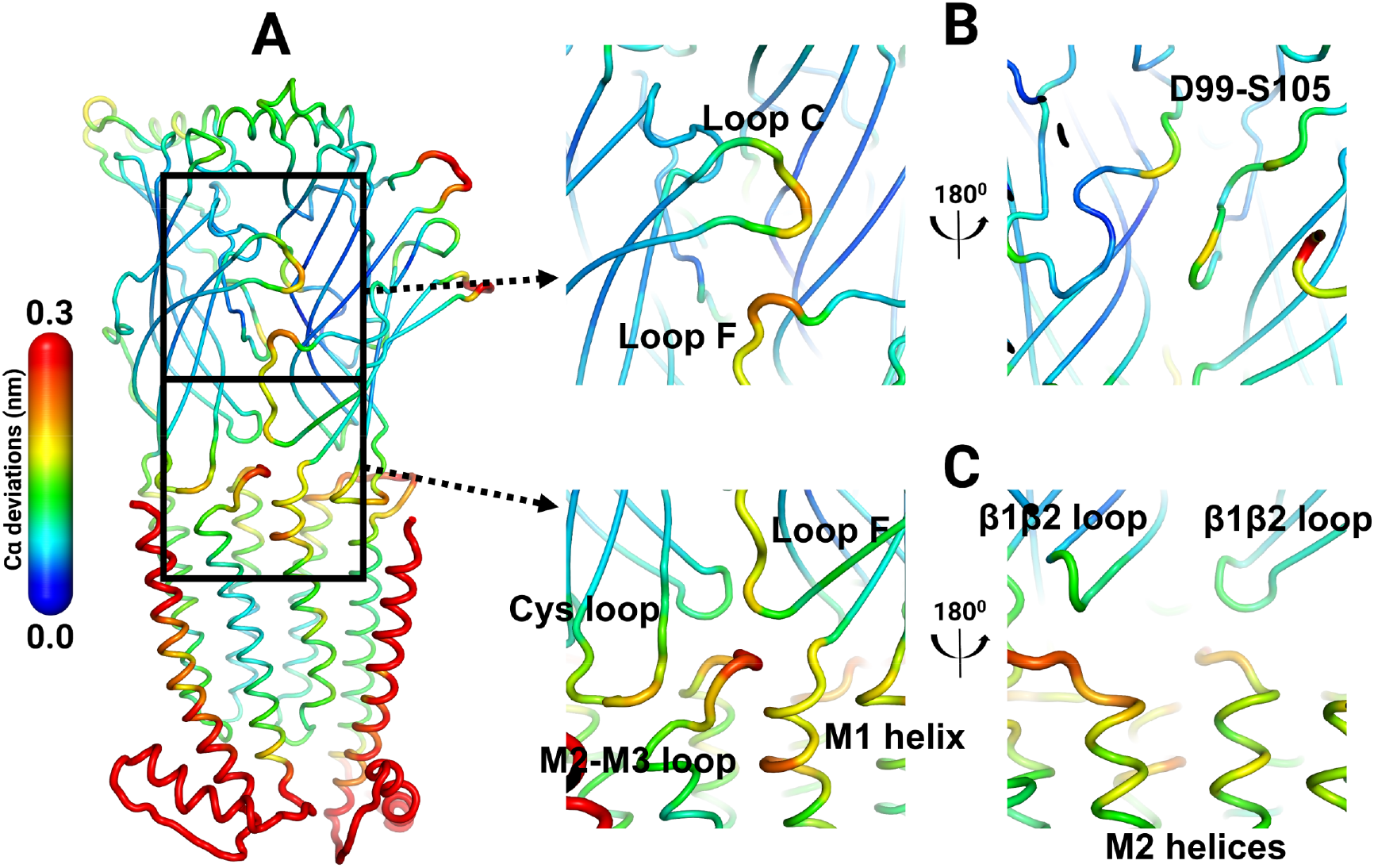
Structural differences between the ACh- and NCT-bound systems. Note that ACh is a full agonist whereas NCT is a partial agonist of the α4β2 nAChR. **A**) Overall structural differences in the α4β2 interface. **B**) Detailed view of the binding pocket. **C)** Detailed view of the ECD/TMD interface. The average Cα positional difference between the ACh- and NCT-bound systems is mapped on the average ACh structure using the colour scale on the left. Note that this figure shows the structural differences in the second α4β2 interface (see Figure S16 for the differences in the first interface).

The comparison between the full and partial agonist bound simulations clearly show that the conformational differences between systems are restricted to the structural motifs involved in receptor activation (*4*), namely the agonist binding pocket, the ECD/TMD interface and the TMD (Figure 2 and S16). Note that caution must be used when assessing the relevance of the structural differences observed in the external regions of the receptor, such as the N- and C-termini and the TM3-TM4 linkers, as these regions are highly flexible. Furthermore, it is also essential to take into consideration that there are missing residues in the termini regions of each subunit and that the ICDs (located between TM3-TM4) were not modelled here, which could lead to an artificial enhancement of the structural differences observed for those regions.

In the orthosteric pocket, structural differences are observed in loops C and the upper part of F and the extracellular selectivity filter (Figure 2B and S16B). The remaining motifs forming the pockets, specifically loops A, B, D and E, show (very) small structural differences between the two systems (Figure 2B and S16B). The extracellular selectivity filter is located in the water-filled extracellular vestibule and mutations in this region are known to influence ion conductance not only in nAChR (e.g. (*58–60*)) but also in other pLGICs (e.g. (*60, 61*)). Loop C is a conserved structural motif that forms the outer face of the binding pocket and it is involved in ligand binding (e.g. (*17*)). Loop C dynamics have been suggested not only to modulate signal propagation (*50, 51*) but also control the mode of action of the agonists. Here, ACh due to its smaller size stabilises more contracted, closed loop C conformations than the bulkier partial agonist NCT (Figure S17).

To identify the associated motions of loop C, the correlation matrix was determined for the ACh- and NCT-bound systems. The correlation values, which can vary between −1 and 1, indicate if two groups tend to move in opposite/similar directions. In our simulations, the motions of loop C in both the ACh- and NCT-bound systems, although tightly coupled to specific regions of the extracellular domains (such as loops A, E and F) show (almost) no correlation with the transmembrane part (Figure S18A), in agreement with mutagenesis studies that have shown that gating can occur without loop C (*17*). Overall, in the binding pocket region only small differences in the correlated motions are observed for the full and partial agonist-bound systems (Figure S18B).

At the ECD/TMD interface, the conformational differences between the systems with partial and full agonists are restricted to three regions, namely M2-M3, Cys and (lower part of) F loops (Figure 2C and S16C). Note that despite variations in amplitude, similar structural rearrangements are observed for the two αβ interfaces. As stated previously, although loop F is essential for binding, its contribution to gating remains elusive (*56, 57*). The concerted motions of loop F were also determined, and the dynamics of its upper part (residues S168-F172) is highly coupled to the movements of the binding site region (Figure S19). The motions of the lower part of the loop (residues P174-W178) are correlated not only with the binding pockets but also with the M2-M3 linker (Figure S20). Interestingly, in the ACh-bound system, the connections of loop F to key functional regions of the receptor, namely loop B, M2-M3 linker and Cys loop, are stronger when compared to the NCT-bound system (Figures S19-S20). This difference in behaviour may suggest a role for loop F in discriminating between partial and full agonists.

The extracellular M2-M3 linker forms part of the ECD/TMD interface and it connects transmembrane helices TM2 and TM3 (*4*). This loop is a well-established gating control element (*62–67*) and mutations here were shown to disrupt the communication between domains and the link between ligand binding and channel activation not only in nAChRs (e.g. (*62–65, 67, 68*)) but also in other pLGICs (e.g. (*69*)). Several computational studies (e.g. (*9, 70*)), including by ourselves (*50, 51*), have emphasised the role of this loop in interdomain signal propagation providing a direct link between the ECD and TMD. Here, the motion of the M2-M3 linker is highly concerted not only with TM1, TM2 and TM3 and the Cys loop region but also with loop F (Figure S21A). The difference in the correlations for the M2-M3 linker between the ACh- and NCT-bound systems is indeed striking (Figure 21B) with the first showing connections spanning all the way through the binding pockets.

The Cys loop is a structural motif common to all pLGIC subunits and the major contributor to the ECD/TMD interface (*4*). This region contains several highly conserved residues, including the canonical FPFD motif (*4*). Site-directed mutagenesis and functional studies have shown this region to be of critical importance for coupling agonist binding to gating (e.g. (*71–74*)) and several disease-causing mutations have been identified here (e.g. (*75*)). In our simulations, the Cys loop dynamics is strongly concerted with the binding pockets, the M2-M3 linker and transmembrane helices TM1, TM2 and TM3 (Figure S22A). Such correlations seem to emphasise the role of this loop as a critical coupling element between domains. Once more, differences in the correlation patterns between the full and partial agonist-bound systems are observed, and they extend all the way to the (+) and (-) side of the binding pockets and to loop F (Figure S22B).

In the ion channel, some structural differences are also observed between the ACh- and NCT-bound systems (Figure S23). However, these differences are restricted to the top region of TM2, mainly between positions 16’ and 20’ (Figure S23). The TM2 helix from each subunit borders the ion pore and mutations can affect ion conductance and cause diseases (*5*). In particular, in the α4β2 subtype, TM2 mutations cause autosomal dominant nocturnal frontal lobe epilepsy (*5*).

The shape of the ion channel was also monitored over the simulation time, and in both systems, the TM2 helices moved from a V-shaped orientation (Figure 3A) to being nearly perpendicular to the membrane (Figure 3B and 3C). At the beginning of the simulations, the TM2 helices had their extracellular ends tilted outward with constriction on the cytoplasmic side in position −1’ due to the side chains of E247 (α4 subunit) and E239 (β2 subunit) (Figure 3A and Figure S24). This closed V-shaped conformation is thought to reflect a desensitised, non-conducting state of the receptor (*7*), which is characterised by high affinity for the agonist and is reached after prolonged or repetitive agonist binding (*4*). However, during the simulation time, a straightening of the TM2 extracellular region was observed leading to the complete closing of the ion channel (Figure 3B, 3C and S24). Despite some variability in the amplitude of the TM2 motions among replicates, similar TM2 rearrangements were observed for the ACh- and NCT-bound systems (Figure S24-S25). Overall, after 280 ns, no significant difference is found in the average diameter of the pore between the systems with full and partial agonists (Figure S26), in agreement with the view that agonist efficacy is independent of the open-close transition (*39*). Several previous computational studies, by ourselves and other groups, have reported similar tilting/straightening motions of the TM2 helices not only in nAChRs (e.g. (*50, 77*)) but also in other pLGICs (e.g. (*11*)). Such motions have been suggested to be essential for ion channel opening and closing. The membrane thickness was also monitored during the simulations (Figure S27-S28), with no significant differences observed between the ACh- and NCT-bound systems.

**Figure 3.**
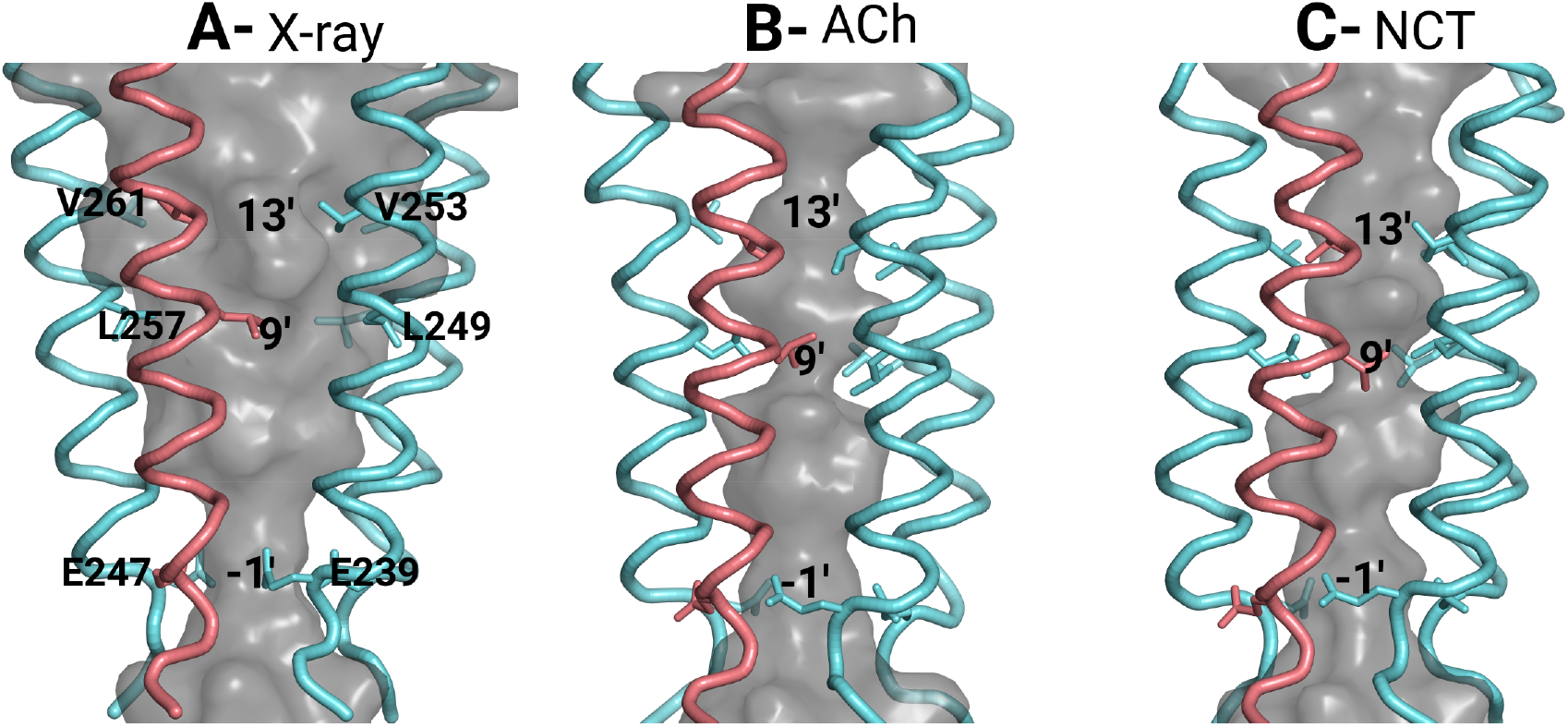
Structural rearrangements in the pore during the simulations. **A**) Ion permeation channel in the X-ray structure used as the starting point for the simulations. **B** and **C**) Ion channel in a representative conformation of the closed channel for the ACh- (**B**) and NCT-bound (**C**) systems. In this image and for simplicity, only the TM2 helices are depicted. The α4 and β2 subunits are coloured in pink and cyan, respectively. Rings of conserved leucines (L257 in the α4 and L249 in the β2 subunits) and valines (V261 in the α4 and V253 in the β2 subunits) located in positions 9’ and 13’ in the TM2 helices form the hydrophobic gate (*4*). The side chains of the residues forming the hydrophobic gate and the cytoplasmic constriction (E247 and E239 in position −1’) are represented with sticks. The grey surface corresponds to the internal surface of the ion channel and was determined using the software HOLE (*76*).

### Signal propagation from the binding pockets to the TMDs

In order to reveal the motifs and the order of the events involved in the propagation of the gating signal generated by ACh and NCT, a large set of complementary D-NEMD simulations were performed. It should be noted that these nonequilibrium simulations should be viewed as complementary to the equilibrium simulations described above, with the former allowing mapping of the agonist-induced conformational changes (after hundreds of nanoseconds), while the latter provides insight into the order of the events associated with signal propagation.

For each agonist-receptor complex simulated, 410 short, 30 ns-long, D-NEMD simulations were performed, adding to a total of 24.6 μs of simulation. All nonequilibrium simulations started from conformations extracted from the equilibrated part of the long equilibrium agonist-bound simulations (Figure S3) and in all of them, the two agonist molecules, either ACh or NCT, were (instantaneously) ablated from the binding pockets forcing signal transmission within the protein (similarly to (*50, 51*)). The Kubo-Onsager approach (*78–81*) was subsequently used to extract the response of the receptor as its conformation adapts to the removal of the agonists and identify the mechanical and dynamic coupling between the structural motifs involved in this response (Figures S29-S32). The response over time was computed by directly comparing the evolution of the simulations with and without agonist bound (similarly to (*50, 51*)). This subtraction approach, and the averaging over hundreds of replicates, allows not only for the temporal sequence of the conformational changes to be identified (Figures S29-S32) but also to assess their statistical significance (Figures S33-S34). At this point, it seems crucial to call the reader’s attention to the fact that due to the artificial nature of the perturbation used here (the instantaneous removal of the agonists), the timescales observed for the response do not correspond to the biological timeframe. Additionally, and due to their short length (30 ns), the nonequilibrium simulations were not intended to model the gating process; instead, they were designed to force signal transmission within the receptor and identify the structural changes associated with signal propagation and the routes by which those rearrangements are transmitted.

The D-NEMD simulations performed clearly identify several common features associated with the structural response to ACh and NCT removal (Figure 4 and S35). Note that for both agonists, interdomain signal transmission starts in the α4 subunit mainly in loop C, and it then propagates to the β2 subunit via loop F and only after that to the TMDs. As can be seen in Figure 4 and S35, 50 ps upon agonist removal, loop C (residues Y197-Y204) is the only region in the binding pockets that undergoes a conformational change. Note that although it is the same region responding, the structural rearrangements are more pronounced in the NCT-bound systems. Over the next 30 ns, an increase in the deviations is observed, propagating from loop C to loop F and afterwards to the TMDs via the M2-M3 linker (Figures 4 and S35).

**Figure 4.**
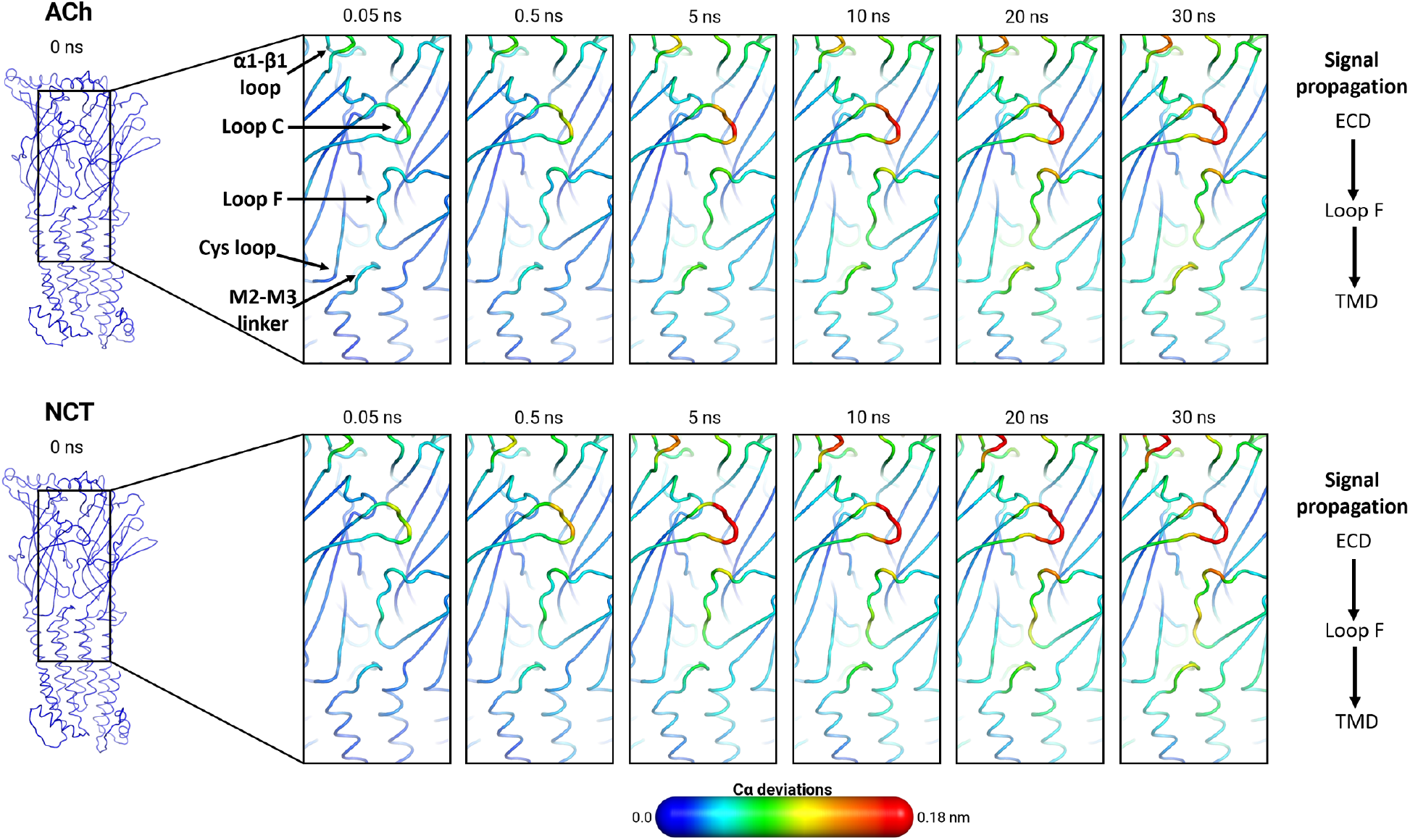
Interdomain signal propagation for the ACh- and NCT-bound systems. The average Cα positional deviation in the 30 ns following agonist removal from the first binding pocket is mapped on the average agonist-free structure. The Cα differences between the nonequilibrium agonist-free and the equilibrium agonist-bound simulations at specific times (0, 0.05, 0.5, 5, 10, 20 and 30 ns) were calculated as a function of the residue number. The final deviation values correspond to the average obtained over all 410 pairs of simulations (Figures S29 and S31).

Nonetheless, and despite using the same pathway for signal propagation to the TMDs, differences in the amplitude of the conformational changes and in rate of transmission of those rearrangements are observed between the ACh- and NCT-bound systems (Figures 5 and S36 and Movie 1). Note for example, that after 30 ns the structural displacements in loops C and F are generally larger in the NCT-bound system.

**Figure 5.**
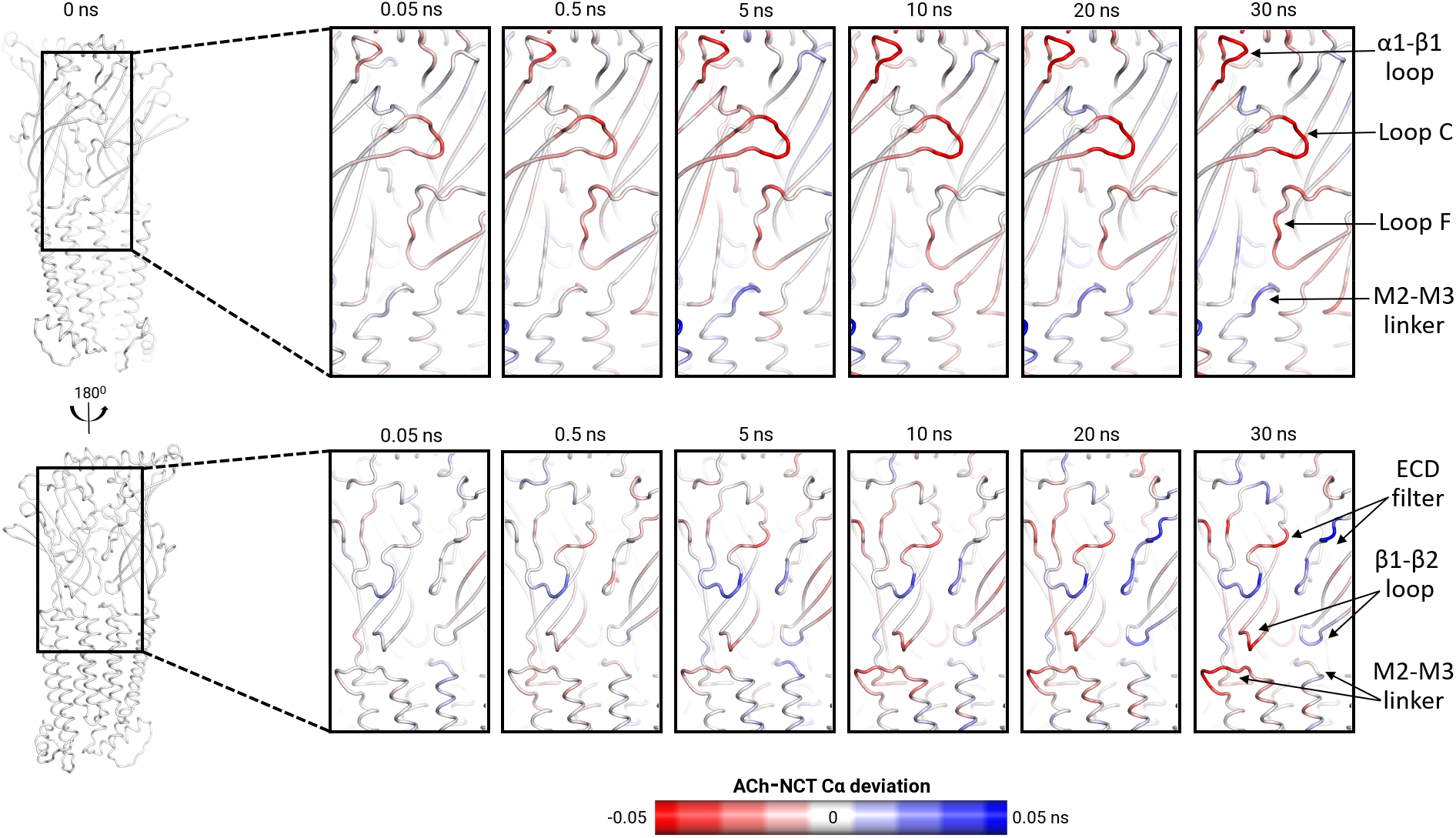
Differences in response to agonist removal between the ACh- and NCT-bound systems. The difference in the average Cα positional deviation between the ACh- and NCT-bound systems in the 30 ns following agonist removal from the first binding pocket is mapped on the average agonist-free structure.

All the structural motifs identified here as being part of the inter-domain communication pathway, namely loops C, F and M2-M3, have been shown experimentally to be involved in ligand binding and/or signal transduction. Although voltage-clamp experiments have demonstrated the role of loop F in ligand affinity, and specificity (e.g. (*82*)), its involvement in gating is still unclear with several works suggesting a role in signal propagation (e.g. (*83, 84*)). Mutations, insertions and deletions of residues in the M2-M3 linker are known to affect ECD/TMD communication and gating (e.g. (*62–65, 67, 68, 85–87*)). Loop C is essential for binding (e.g. (*17, 88*)) by stabilising the positively charged group of the agonists; however, no consensus exists regarding its role in the mode of action of the agonists.

Additionally, as suggested by us previously (36) and contrary to other motifs (e.g. loop C), the Cys loop responds very slowly to agonist removal. It takes several tens of nanoseconds to start undergoing any structural rearrangements (Figures 4 and S35). However, after 30 ns, no difference in behaviour is observed for the Cys loop between the systems with full and partial agonists (Figures 5 and 36).

Remarkably, the α1-β1 loop, which is located above the binding pocket in a peripheral region of the receptor, also responds rapidly to agonist removal (Figures 4 and S35) and shows an agonist-specific response with larger structural rearrangements in the NCT-bound system (Figures 5 and S36). These conformational changes although not relevant for inter-domain signal propagation (as they are located far from the ECD/TMD interface) may be necessary for interactions with other proteins and modulators (for a review see, e.g. (*89*)).

Intriguingly, the residues located in the extracellular selectivity filter (*60*) also react to agonist removal (Figures S37-S38). After 30 ns, differences in the amplitude of the conformational changes between the full and partial agonist-bound systems are observed (Figures 5 and S36). While ACh removal induces larger rearrangements in the loop forming the ECD filter from the α4 subunit, NCT removal affects mostly the loop from β2 subunit (Figures 5 and S36). The ECD filter has a role in ion discrimination, thus enhancing the receptor’s selectivity for small cations (e.g. Na^+^ and K^+^) (*21*). Mutations in the ECD vestibule were shown to influence ion conductance (e.g. (*58, 59, 61*)).

Finally, but importantly, the nonequilibrium simulations performed also allow us, for the first time, to directly follow the structural response of the receptor as it propagates through the ion channel (Figures S39-S40 and Movie 2). Despite no significant differences observed between the ACh- and NCT-bound systems (within the simulated timescale), it is still fascinating to follow the order of the events associated with signal propagation along the TM2 helices. Unsurprisingly, the conformational changes inside the pore start in the extracellular end of TM2 helix (around position 20’), and over time, they propagate downwards in a wave-like manner, reaching the hydrophobic gate (between position 9’ and 13’) after 30 ns (Figure S39-S40). Note the time lag (of about 0.5 ns) between the structural rearrangements starting in the ECD and those in the ion pore (Movie 2).

## Concluding remarks

Despite decades of intensive study, the structural mechanisms underlying agonist efficacy remain poorly understood. A detailed description of the structural changes induced by full and partial agonists and how these rearrangements propagate within the receptor is fundamentally important for a molecular understanding of each agonist’s mode of action. In this work, we made use of a combination of equilibrium and nonequilibrium MD simulations to study the behaviour of the human α4β2 receptor with two agonists bound, namely acetylcholine (full agonist) and nicotine (partial agonist) and identify the differences in their working mechanisms. Our equilibrium simulations show that the ligands induce different structural and dynamic changes in the receptor. The largest difference in dynamics between the two agonist-bound systems occurred in loop F, a motif known to be involved in binding affinity, specificity and possibly gating. The structural differences between the full and partial agonist-bound systems were restricted to loop C, the ECD selectivity filter and the ECD/TMD interface, notably in the M2-M3, Cys and F loops.

The nonequilibrium simulations not only identify the pattern of through-receptor communication and the order of the conformational changes associated with signal transmission but also allows for the mapping of the progress of the agonist-specific structural rearrangements over time. These structural rearrangements always start in the binding pockets with subsequent transmission to different parts of the receptor. Although ACh and NCT use the same pathways for signal transmission, differences in the amplitudes of the conformational response of key functional motifs and rates of propagation are observed between the agonists. Overall, over the 30 ns, ACh triggers larger changes in the loop β1-β2 and ECD filter from the α4 subunit while NCT steadily induces larger rearrangements in loops C and α1-β1 from the α4 subunit and loops F, β1-β2 and ECD filter from the β2 subunit. These conformational and temporal differences relate to agonist structure and may correlate efficacy to the diverse structural response of the α4β2 nAChRs to a ligand.

## Materials and methods

Molecular dynamics (MD) simulations of the human (α4)_2_(β2)_3_ nAChR (PDB code: 5KXI) (*7*) with two agonists bound, namely acetylcholine (full agonist) and nicotine (partial agonist), were carried out to identify the agonist-induced conformational changes and how the ligands modulate the dynamics of the receptor. For each complex, ten replicates, 280 ns long (totalling 2.8 μs per system) were performed with the receptor inserted into an explicit lipid membrane (for details, see the supplementary information). Additionally, an extensive complementary set of 820 (410 for the ACh- and 410 for the NCT-bound systems), 30 ns-long, dynamical nonequilibrium MD (D-NEMD) simulations (in a total of 24.6 μs) was acquired to study how the agonists modulate signal propagation (see supplementary information for a detailed description). Hundreds of simulations were required to achieve convergence and demonstrate statistical significance. In all nonequilibrium simulations, the agonists were removed from the binding pockets as in (*50, 51*), and the trajectory of the resulting agonist-free system was followed for 30 ns. These simulations were designed to drive and allow for the identification of the conformational effects of ligand removal, the coupling between structural elements, and the order of the events associated with signal propagation. The characterisation of the temporal evolution of conformational changes in the system as a response to agonist removal was extracted using the Kubo-Onsager approach (*78–80*). All simulations were done with Gromacs (*90–92*) on the University of Bristol’s High-Performance Computer, BlueCrystal. All simulations conditions were the same as described in (*50, 51*).

## Acknowledgements

AJM and ASFO thank EPSRC (grant number EP/M022609/1), BBSRC (grant number BB/R016445/1) and ERC (grant PREDACTED https://cordis.europa.eu/project/id/101021207)for support. We also thank BrisSynBio, a BBSRC/EPSRC Synthetic Biology Research Centre (Grant Number:BB/L01386X/1) for funding ASFO. MD simulations were carried out using the computational facilities of the Advanced Computing Research Centre, University of Bristol (http://www.bris.ac.uk/acrc).

## Data access statement

The data is available upon request.

