## Supplementary Information for "Structural and temporal basis for agonism in the α4β2 nicotinic acetylcholine receptor"

#### Starting Structure

The 3.9 Å X-ray structure of the human $(\alpha4)_2(\beta2)_3$ nicotinic acetylcholine receptor (nAChR) with nicotine bound (PDB code: 5KXI) (*1*) was used as the starting point for this work. All the missing loops were modelled with the program MODELLER 9v17 (*2, 3*), similarly to (*4*). It should be noted that in this work, the intracellular domain of the receptor was not modelled due to the lack of structural information.

For the system with acetylcholine bound, acetylcholine was placed in the binding pockets in positions analogous to the ones occupied in the acetylcholine binding protein's crystal structures from *Aplysia californica* in complex with acetylcholine (*5*).

#### Insertion of the receptor into a lipid bilayer

Two systems were prepared, namely an acetylcholine-bound (ACh) and a nicotine-bound (NCT) system (Figure S1). The agonist-receptor complexes were inserted in a pre-equilibrated POPC bilayer using LAMBADA and InflateGRO2 (*6*).

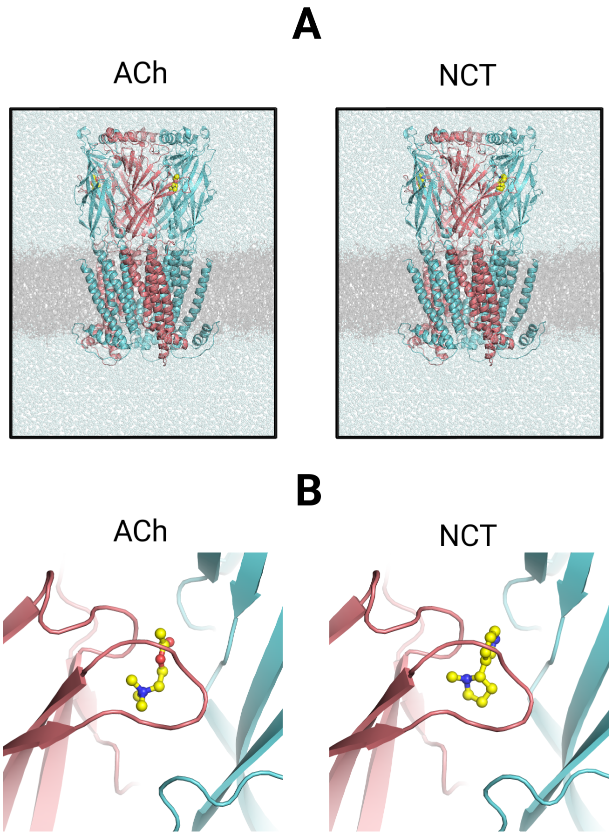

**Figure S1. A)** View of the ACh- and NCT-bound systems. The protein is rendered as a cartoon with the α4 subunits coloured in pink and the β2 in cyan. The lipids are represented as grey sticks. The agonists (ACh in the left-side image and NCT in the right-side image) are shown with yellow spheres. **B)** Initial binding mode of ACh and NCT inside the binding pockets.

ACh is a full agonist of the human α4β2 nAChRs, meaning that it produces maximum responses (*7*). On the other hand, NCT elicits less than the maximum response (even at maximally effective concentrations) when compared to the ACh (*8*), and, hence, is a partial agonist of the α4β2 receptors (Figure S2). Note that efficacy is not an inherent property of the agonist and it is specific for different agonist-subtype complexes, with several examples of molecules acting as full agonists in one nAChR subtype and partial agonists in another (*7*). In this work, we have focused on the human α4β2 nAChR for clinical reasons as this subtype is the most prevalent nAChR in the brain with high affinity for nicotine (*9*) and, thus is key for nicotine addiction.

The two agonist-receptor-membrane complexes were solvated using TIP3P water molecules (*10*). An ionic concentration of 0.1 M sodium chloride was used similarly to (*4*). The total system size was ~275K atoms.

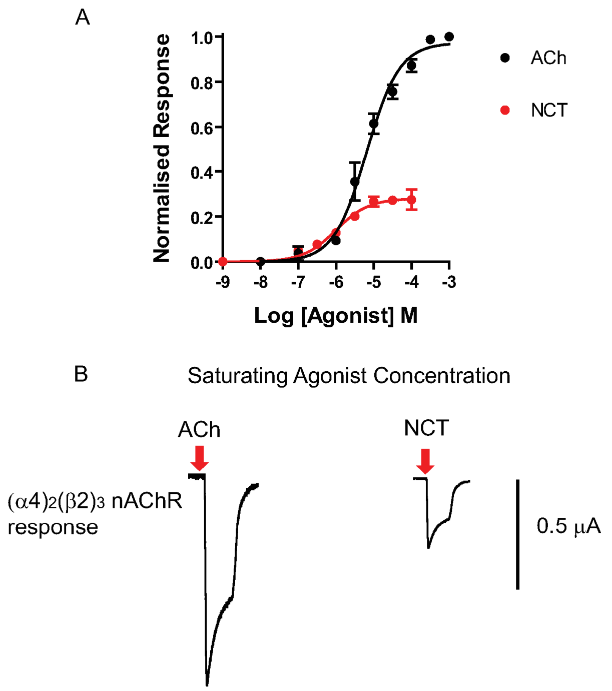

**Figure S2.** Maximal responses of $(\alpha4)_2(\beta2)_3$ nAChR to ACh and NCT. **A)** Concentration-response curves for ACh (black dots and curve) or NCT (red dots and curve at $(\alpha4)_2(\beta2)_3$ nAChR. ACh is a full agonist, whereas NCT behaves as a partial agonist, eliciting only approx. 30% of the maximal response elicited by ACh. **B)** Comparison of the amplitude of the maximal currents responses elicited by ACh or NCT at $(\alpha4)_2(\beta2)_3$ nAChR. Data in **A** and **B** were obtained by two voltage-clamp recordings from *Xenopus* oocytes expressing $(\alpha4)_2(\beta2)_3$ nAChRs. Data points in the concentration-response curves shown in **A** represent the mean ± SEM of 10-12 independent experiments carried out using 5 different batches of

Xenopus oocytes. Peak current amplitudes for the two agonists were normalised to 1 mM, a maximally efficacious ACh concentration at the $(\alpha4)_2(\beta2)_3$ nAChRs.

#### Equilibrium MD simulations

All equilibrium MD simulations were performed using Gromacs (version 5.1.4) (*11-13*) on the University of Bristol's High-Performance Computer, BlueCrystal (Phase 4). The Amber ff99SB-ILDN (*14*) and the S-lipids (*15, 16*) forcefields were used to describe the protein and membrane, respectively. The parameters for NCT and ACh were taken from our previously published works (*17, 18*). The agonists were considered to be positively charged. All simulation conditions were similar to (*4*). A time step of 2 fs was used for integrating the equations of motion. Non-bonded long-range electrostatic interactions were calculated using the smooth particle mesh Ewald method (*19*). A 12 Angstrom cut-off was used for the van der Waals interactions with long-range dispersion corrections for the energy and pressure (*20*).

The solvated protein-membrane systems were energy minimised, equilibrated and simulated according to the protocol described in (*4*). Ten unrestrained MD simulations, each 280-ns long, were performed for each system.

##### Dynamical-nonequilibrium MD (D-NEMD) simulations

In order to study signal propagation within the receptor, a large number (410 per system) of short (30 ns) D-NEMD simulations were performed. These simulations drive and allow for the characterisation of conformational changes happening over time by using the Kubo-Onsager approach (*21-24*). The protocol for preparing the short simulations is described in detail in (*4*). Briefly, all nonequilibrium simulations started from conformations extracted from the long equilibrium agonist-bound simulations (Figure S3). From the equilibrated part of the 280-ns agonist-bound simulations (from 50-280 ns, respectively), conformations were extracted every 5 ns and used as starting points for the nonequilibrium simulations (in a total of 41 conformations per replicate). A large number of nonequilibrium simulations is required to achieve convergence and to demonstrate the statistical significance of the structural changes identified. In all extracted conformations, the two agonist molecules were removed from the binding pockets in a procedure similar to (*4, 25*), and the trajectory of the resulting system was followed for 30 ns. Note that the main purpose for agonist removal is to create a perturbation in the system and force a response from the receptor. Additionally, due to their short timescales and artificial nature of the perturbation introduced, the nonequilibrium simulations are not attempting to model gating (*4, 25*).

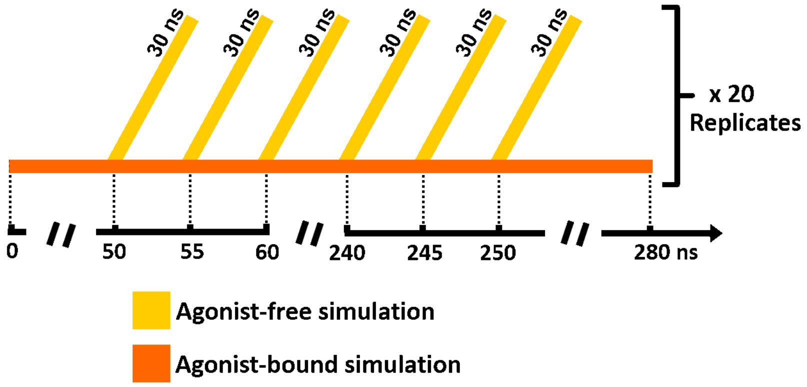

**Figure S3.** Scheme for the nonequilibrium simulations. 20 equilibrium MD simulations (10 for the ACh- and 10 for NCT-bound systems) of the human $(\alpha4)_2(\beta2)_3$ nAChR with agonists bound were performed. These agonist-bound simulations (orange simulations in the scheme) were used to generate starting conformations for the 30 ns-long agonist-free nonequilibrium simulations (yellow simulations in the scheme).

The Kubo-Onsager approach (*21-24*) was used to analyse the nonequilibrium simulations and determine the response of the system to ACh and NCT removal (similarly to (*4, 25*)). According to this approach (*21-24*), the response of a system to a perturbation can be directly measured by averaging a given property in perturbed and unperturbed simulations at a given time, as long as enough statistics are gathered. For each pair of unperturbed agonist-bound equilibrium and perturbed agonist-free nonequilibrium simulations, the difference in position for each Cα atoms was determined at equivalent points in time, namely after 0, 0.05, 0.5, 5, 10, 20 and 30 ns of simulation. The positional deviations obtained at each point in time were, then, averaged over all replicates.

#### Analysis

The RMSD and RMSF were calculated after least-squares fit to the Cα atoms of the initial structure. The DSSP software (*26*) was used to assign secondary structure. PCA was performed to examine the sampling and equilibration of the replicates (similarly to, e.g. (*27-29*)). All replicates for each system were combined before the analysis so that all shared the same subspace and could directly be compared. Each PCA trajectory contained one conformation per nanosecond per replicate. The two principal components (PC1 and PC2) were used to assess the equilibration/relaxation of the simulations, and all systems were considered equilibrated after 50 ns. The Kubo-Onsager approach (*21-24*) was used to analyse the nonequilibrium simulations and determine the response of the system to the removal of the agonists.

#### Conformational stability of the $(\alpha 4)_2(\beta 2)_3$ receptor

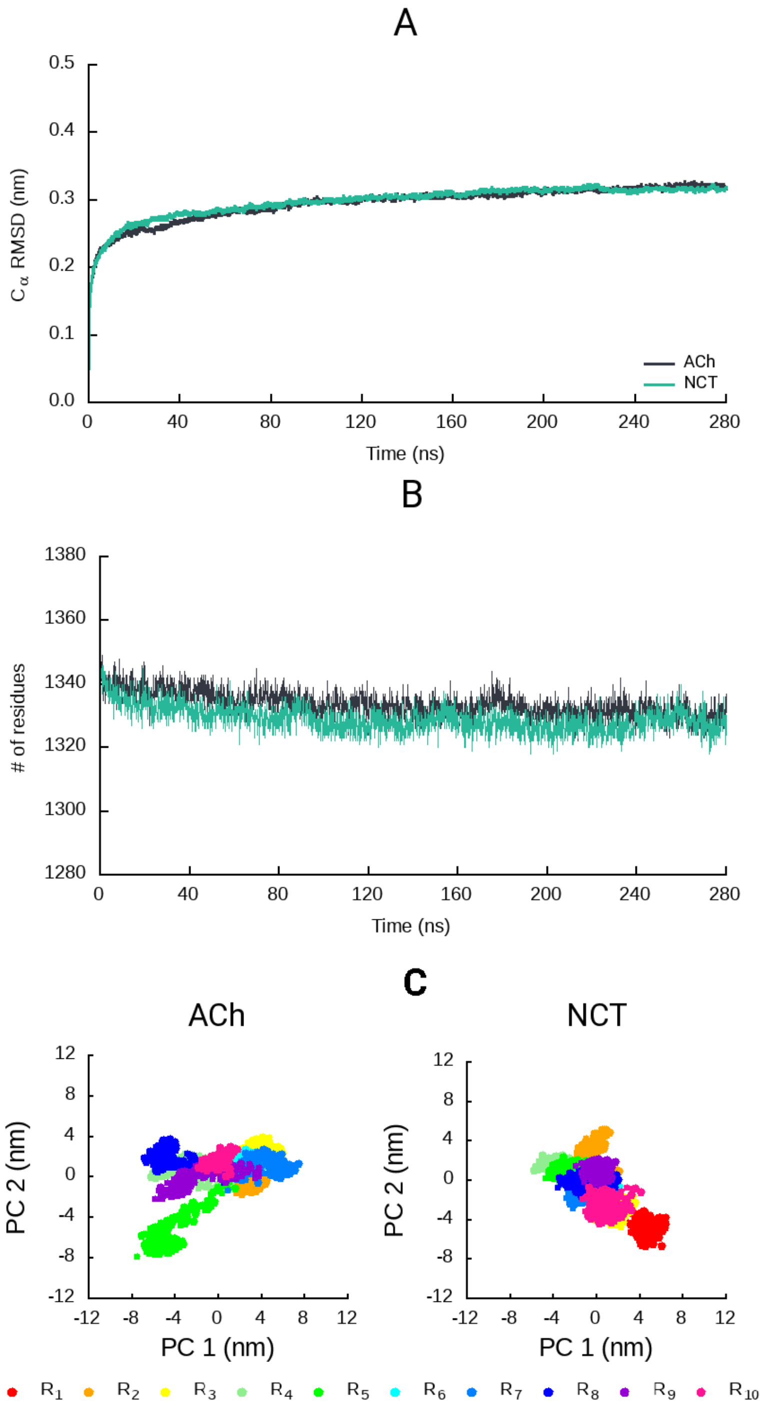

**Figure S4.** Equilibration and stability of the ACh- and NCT-bound systems. **A)** Temporal evolution of the average Cα Root Mean Square Deviation (RMSD) for the ACh- (black) and NCT-bound (green) systems. The Cα RMSD was calculated relative to the starting structures, and the averages were obtained over all 10 replicates. **B)** The number of residues with secondary structure in the ACh- (black) and NCT-bound (green) systems. The DSSP software (*26*) was used for the secondary structure assignment and included all residues assigned to α-helix, $3_{10}$- helix, 5-helix, β-sheet and β-bridge secondary structure classes. **C)** PCA of the ACh- and NCT-bound systems. All replicates were combined before the analysis, and each PCA trajectory contained one conformation per nanosecond per replicate (totalling 5601 frames) with all the Cα atoms of the protein. For a detailed view, zoom into the image.

#### Dynamic behaviour of the agonists during the simulations

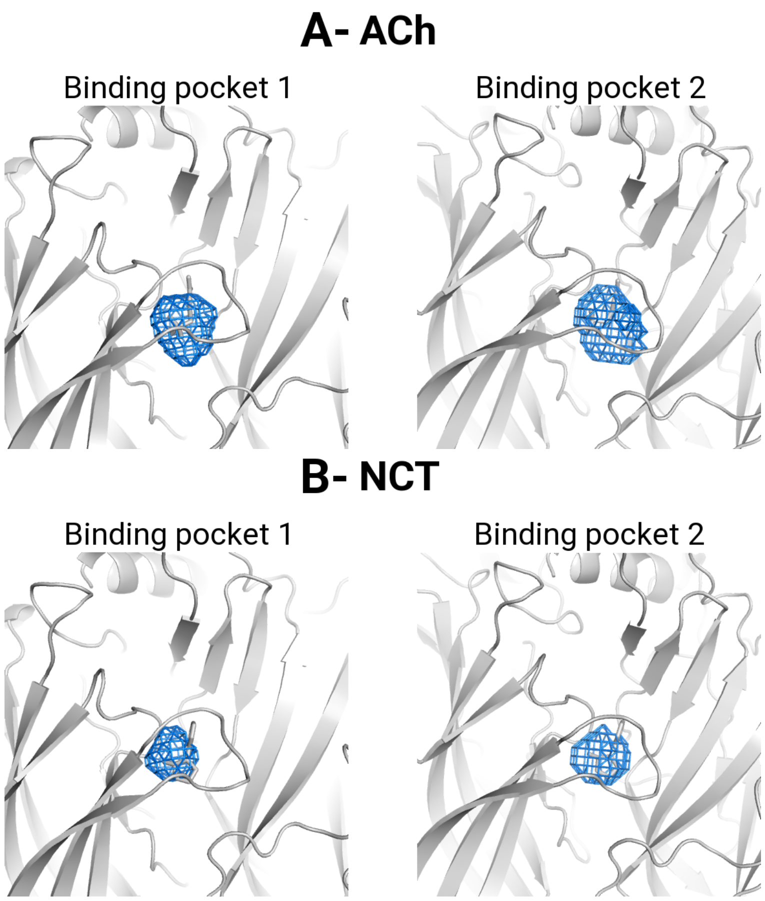

**Figure S5.** Probability density maps for the positively charged group of ACh (**A**) and NCT (**B**). The contours at 0.00001 $Å^{-3}$ are depicted as a blue mesh. The structure used as the starting point for the simulations is shown in grey.

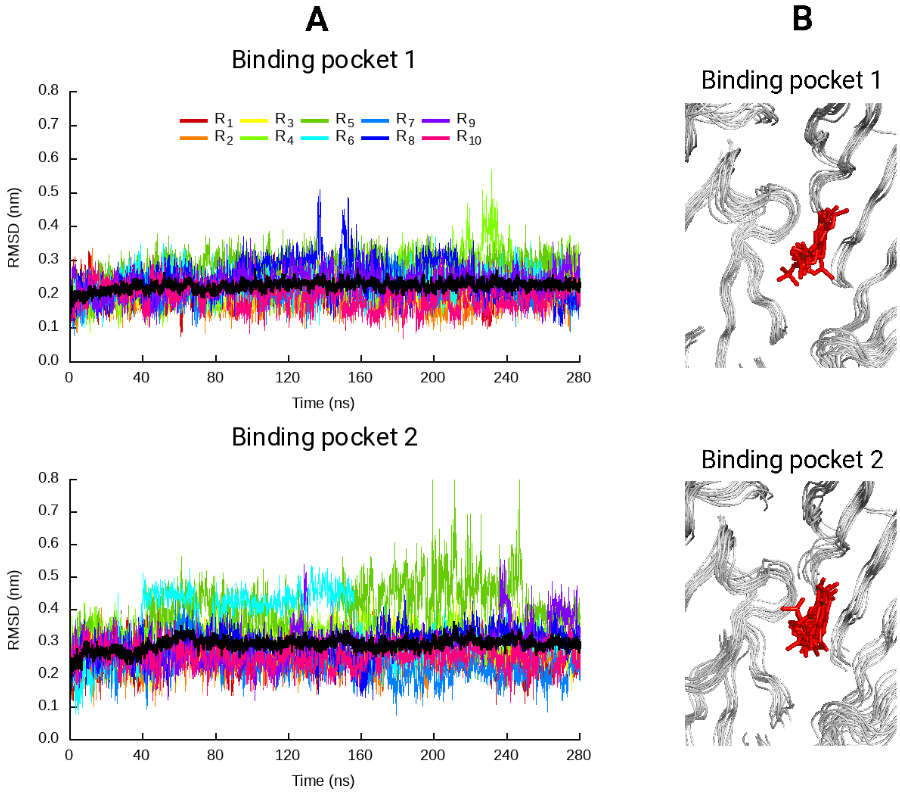

**Figure S6. (A)** Temporal evolution of RMSD of ACh relative to its initial position. The RMSD values were determined considering all the atoms of the agonist. The averages represented (black line) were obtained over all 10 replicates. **(B)** Snapshot of the last frame from all replicates highlighting the binding mode of acetylcholine. Acetylcholine is represented with red sticks.

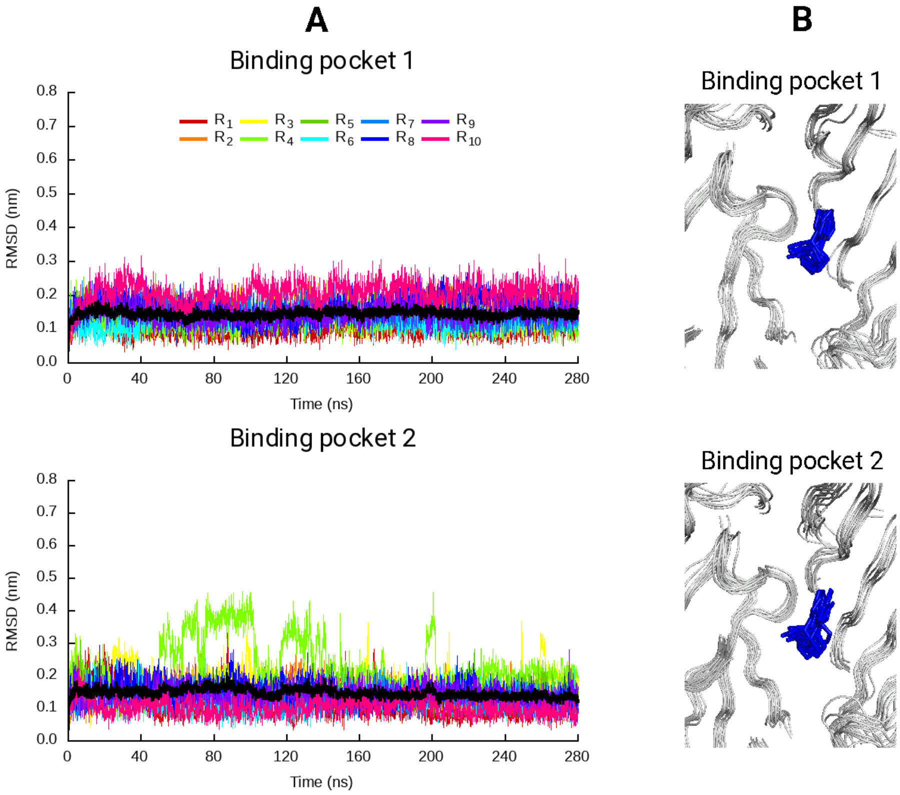

**Figure S7. (A)** Temporal evolution of RMSD of NCT relative to its initial position. The RMSD values were determined considering all the atoms of the agonist. The black line is the average over all 10 replicates. **(B)** Snapshot of the last frame from all replicates highlighting the binding mode of nicotine. Nicotine is represented with blue sticks.

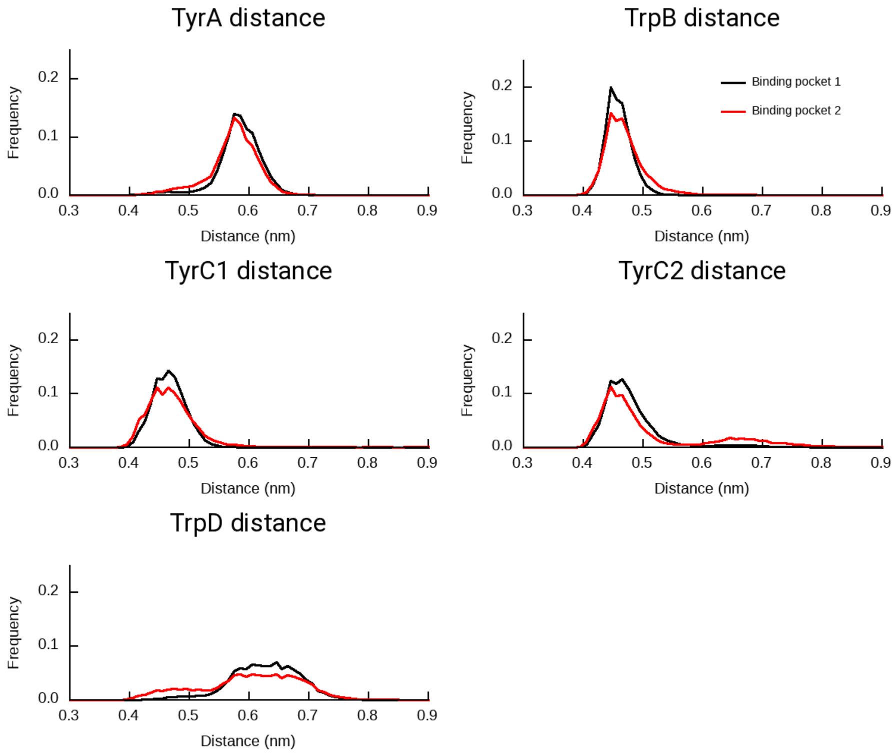

**Figure S8.** Distance between the side-chain of the residues forming the binding pockets, namely TyrA (Y100 in the α4 subunit), TrpB (W156 in the α4 subunit), TyrC1 (Y197 in the α4 subunit), TyrC2 (Y204 in the α4 subunit) and TrpD (W57 in the β2 subunit), and the charged N atom of acetylcholine. The black and red lines correspond to the distance distributions for the first and second binding pockets, respectively.

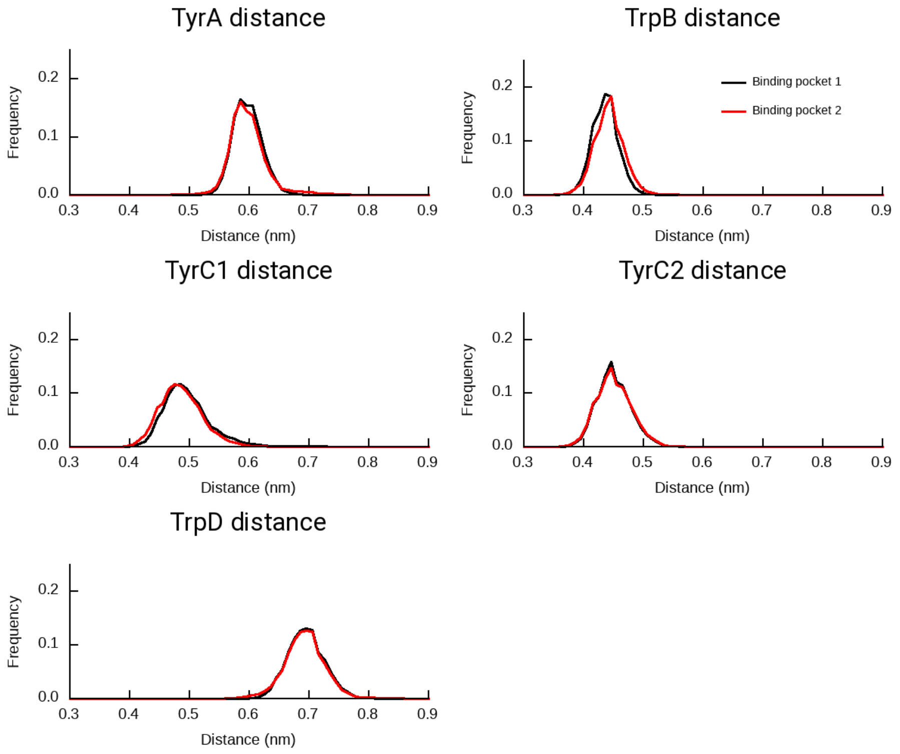

**Figure S9.** Distance between the side-chain of the residues forming the binding pockets, namely TyrA (Y100 in the α4 subunit), TrpB (W156 in the α4 subunit), TyrC1 (Y197 in the α4 subunit), TyrC2 (Y204 in the α4 subunit) and TrpD (W57 in the β2 subunit), and the charged secondary amine N atom of nicotine. The black and red lines correspond to the distance distributions for the first and second binding pockets, respectively.

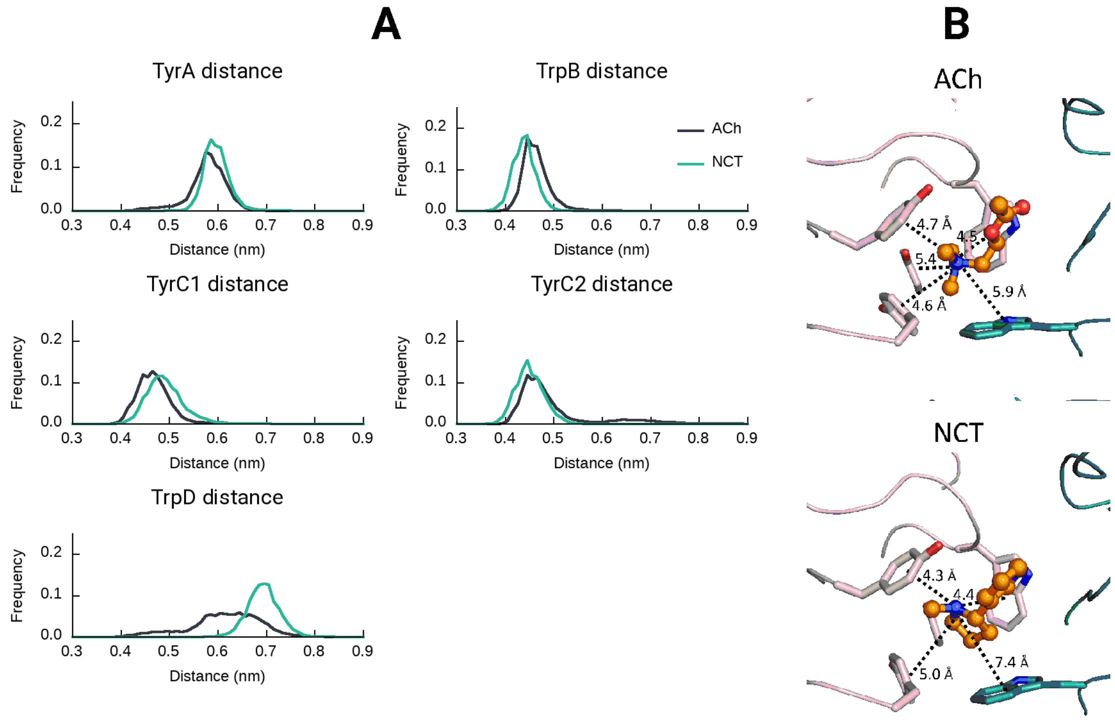

**Figure S10. (A)** Distance between the side-chain of the residues forming the binding pockets, namely TyrA (Y100 in the α4 subunit), TrpB (W156 in the α4 subunit), TyrC1 (Y197 in the α4 subunit), TyrC2 (Y204 in the α4 subunit) and TrpD (W57 in the β2 subunit), and the charged N atom of the agonists. The black and green lines correspond to the distance distributions for ACh and NCT, respectively. These histograms reflect the distances over the two binding pockets. **(B)** Example of the agonists' binding mode inside the aromatic box. The principal and complementary subunits are coloured in pink and green, respectively. The interactions between the positively charged group of the agonists and the aromatic rings of TyrA, TrpB, TyrC1, TyrC2 and TrpD are highlighted with dashed lines.

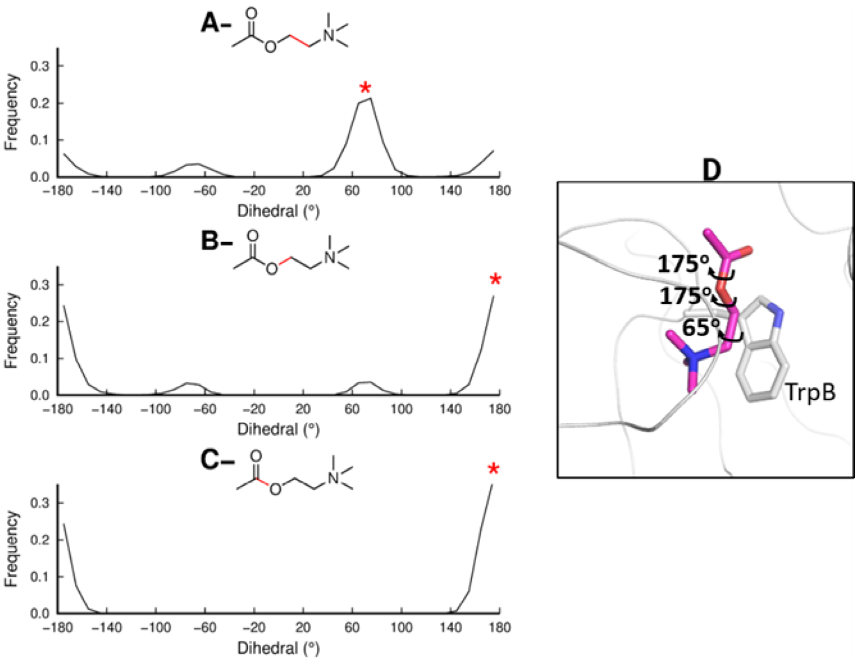

**Figure S11.** Dihedral angles distribution for ACh in the equilibrium MD simulations (**A-C**). These histograms reflect the dihedral distributions in the two binding pockets. The dihedrals measured are highlighted with a red bond. The asterisk corresponds to the most populated angle. **D)** A representative example of the preferred binding mode of ACh inside of the orthostatic pockets.

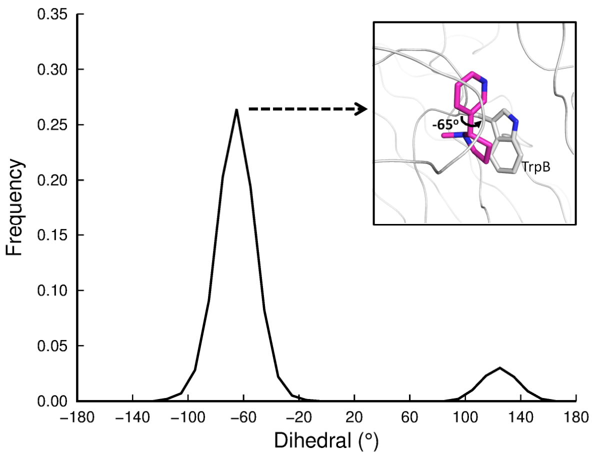

**Figure S12.** Dihedral (between the pyridine and pyrrolidine rings) angle distribution for nicotine in equilibrium MD simulations. This histogram reflects the dihedral distribution for NCT in the two binding pockets.

### Agonist-induced dynamic changes

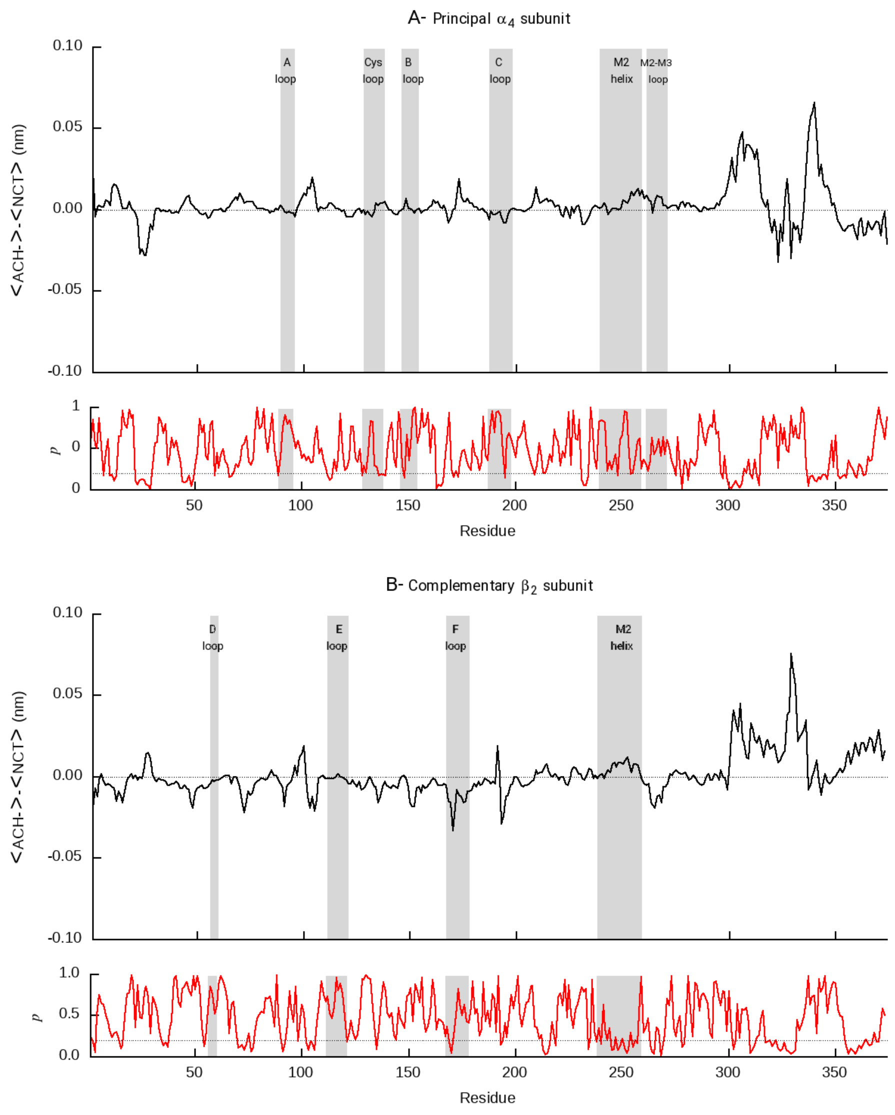

**Figure S13.** Average difference in RMSF between ACh- and NCT-bound systems and associated *p*-values for the principal (α4) and complementary (β2) subunits forming the first binding pocket. The positions of some structural motifs are highlighted in grey. For a detailed view, please zoom into the image.

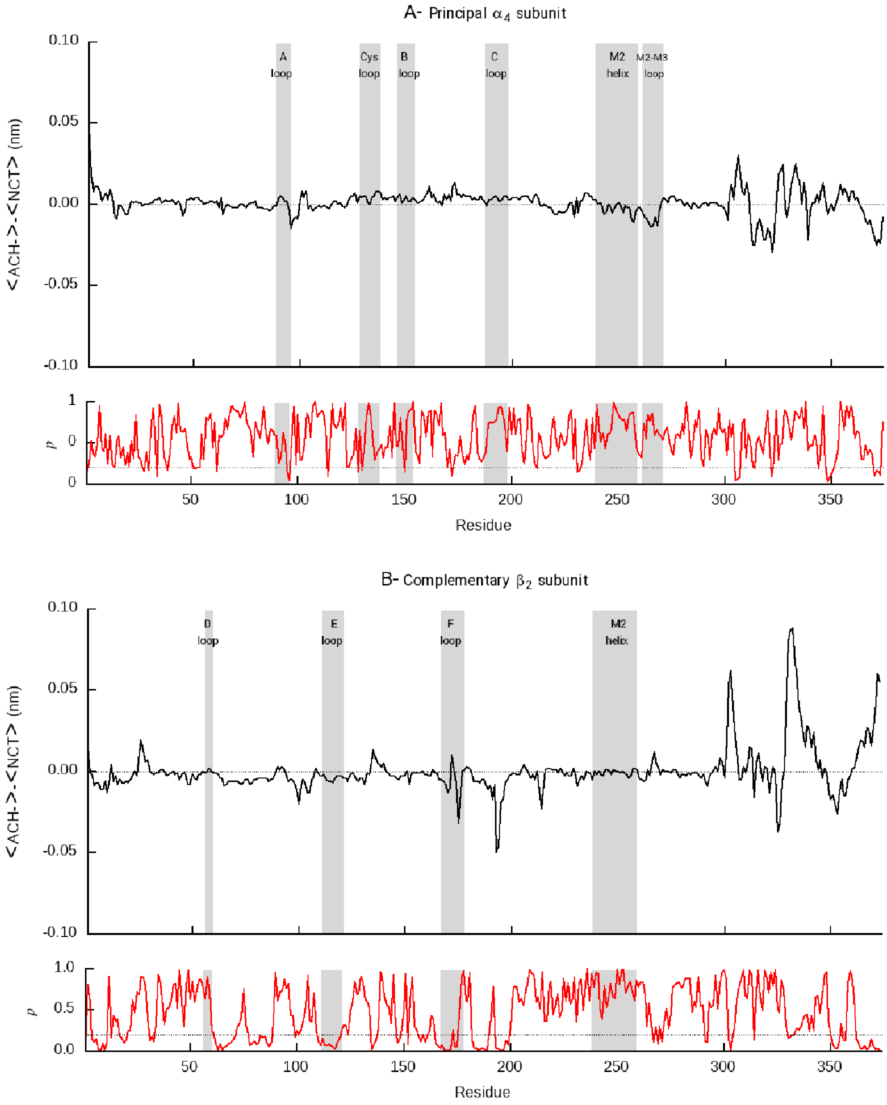

**Figure S14.** Average difference in RMSF between ACh- and NCT-bound systems and associated *p*-values for the principal (α4) and complementary (β2) subunits forming the second binding pocket. The positions of some structural motifs are highlighted in grey. For a detailed view, please zoom into the image.

#### Agonist-induced conformational changes

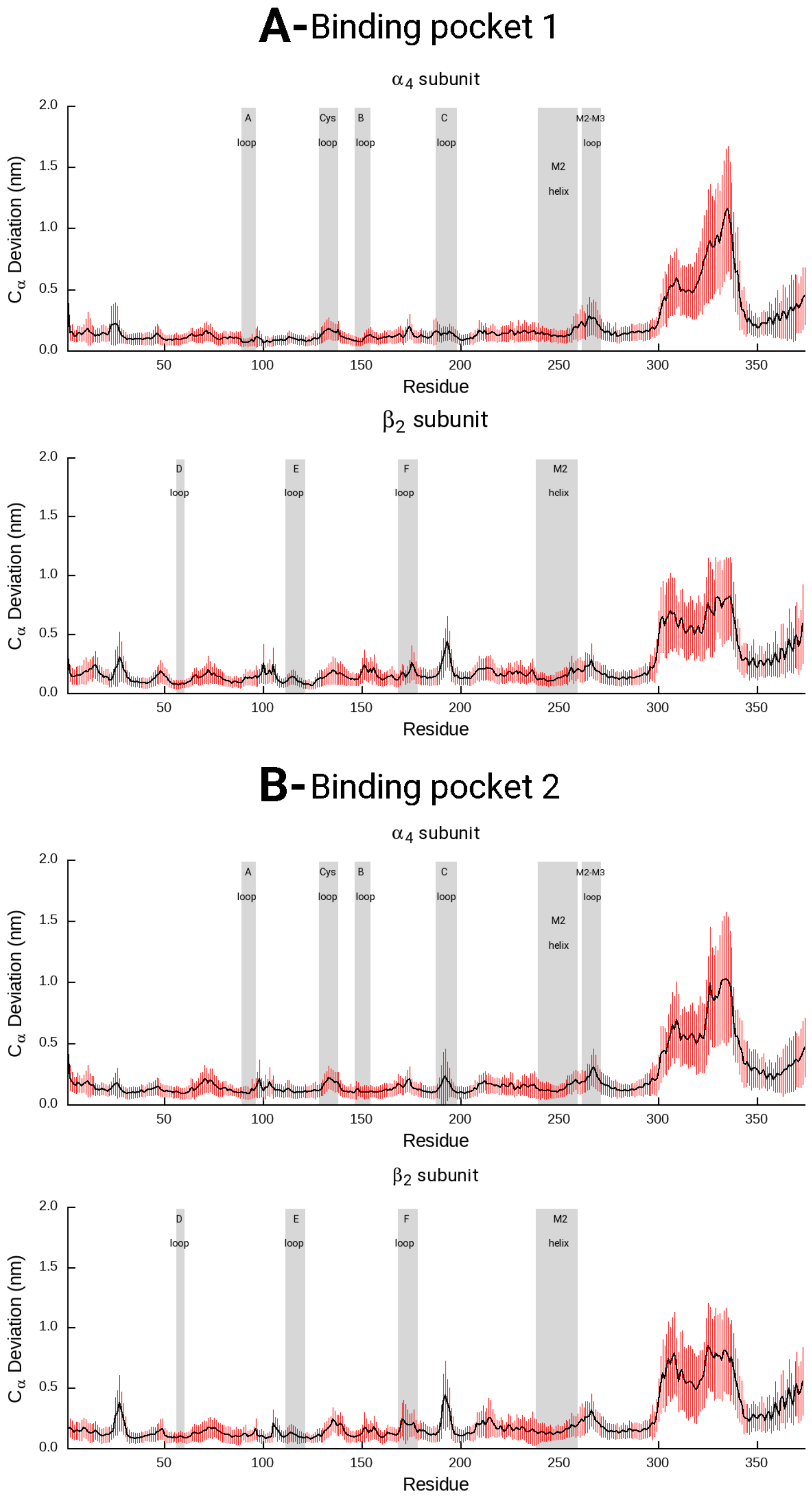

**Figure S15.** Average Cα-positional deviation between the ACh- and NCT-bound systems. The average deviation was determined from all 100 combinations (resulting from the 10 NCT × 10 ACh pairs of trajectories) of $C_\alpha$ RMSD between the average structures of the two systems. The vertical red lines represent the standard deviation of the mean. For a detailed view, please zoom into the image.

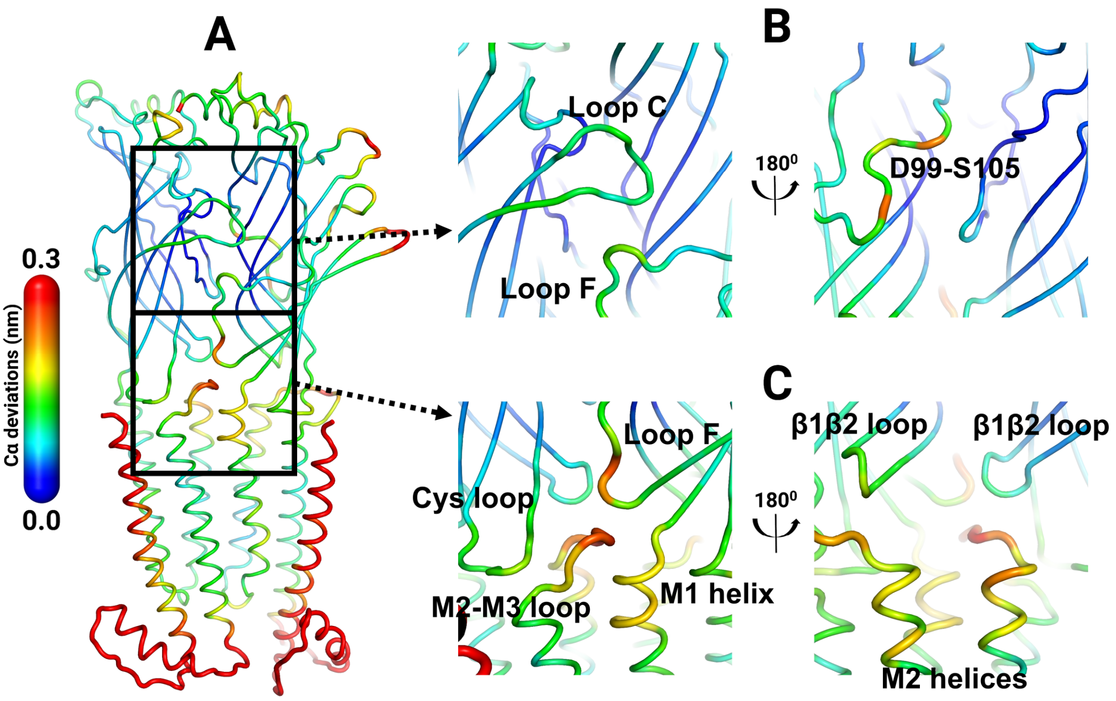

**Figure S16.** Structural differences in the first α4β2 interface. **A**) Comparison between the ACh- and NCT-bound systems. **B)** Detailed view of binding pocket 1. **C)** Detailed view of the ECD/TMD interface. The average $C_\alpha$ positional deviation between the ACh- and NCT-bound systems is mapped on the average ACh structure.

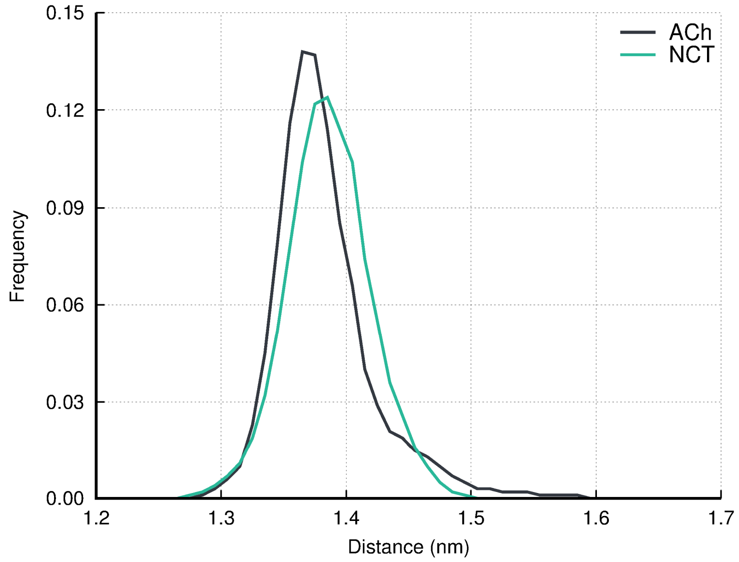

**Figure S17.** Loop C closing in the ACh- and NCT-bound systems. Distributions of the distance between the centre of mass of loop C and the centre of mass of the residues K152 and F153 of the same α4 subunit for the ACh- and NCT-bound systems. The histograms contain the distances for both binding pockets.

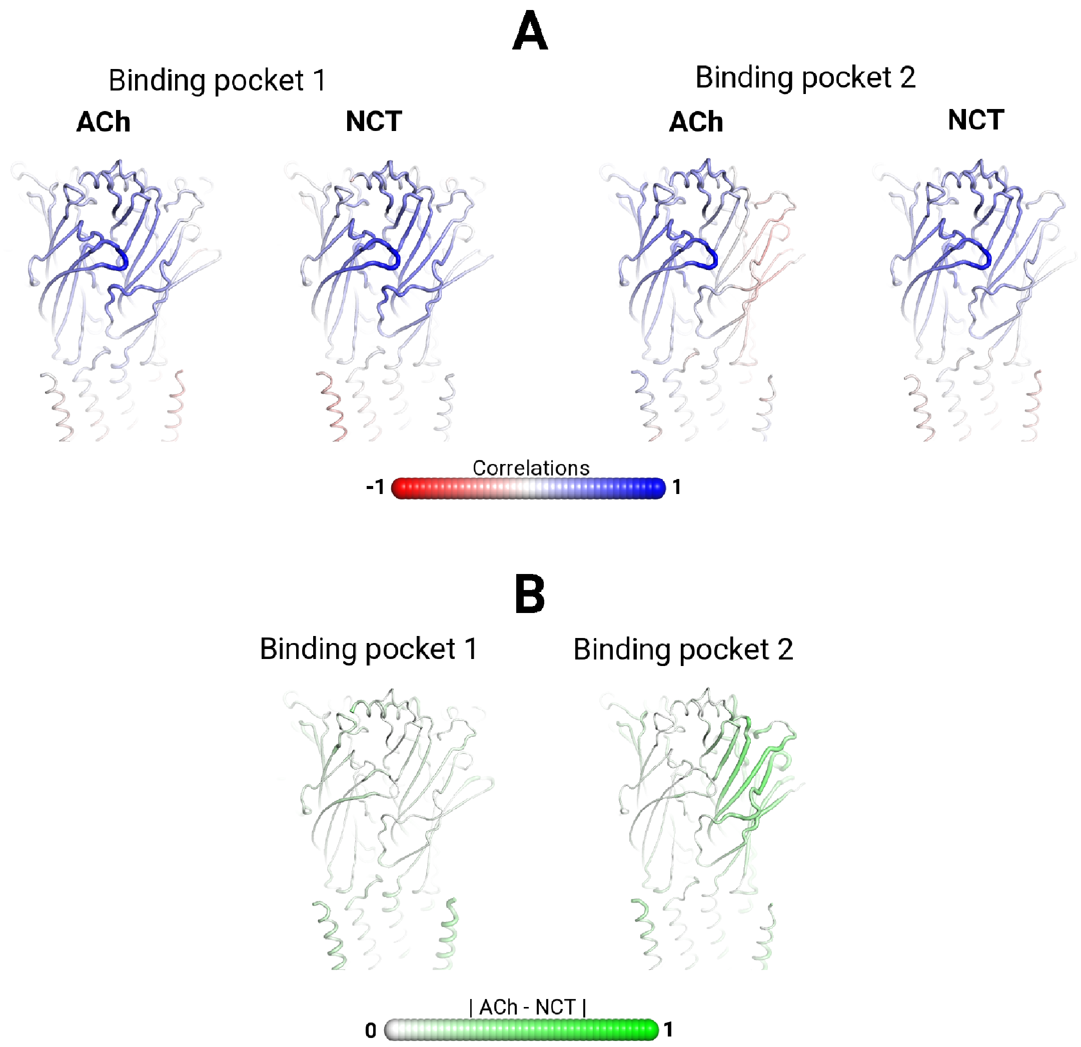

**Figure S18.** (**A**) Statistical correlation between the Cα atom of C199 (located in loop C) in the α4 subunit and all the remaining Cα atoms for the ACh- and NCT-bound systems. In this panel, the atoms that systematically move along the same/opposite direction have a correlation value of 1/-1, whereas those whose movements are uncorrelated present a correlation value of 0. (**B**) Difference between the correlations for the ACh- and NCT-bound systems.

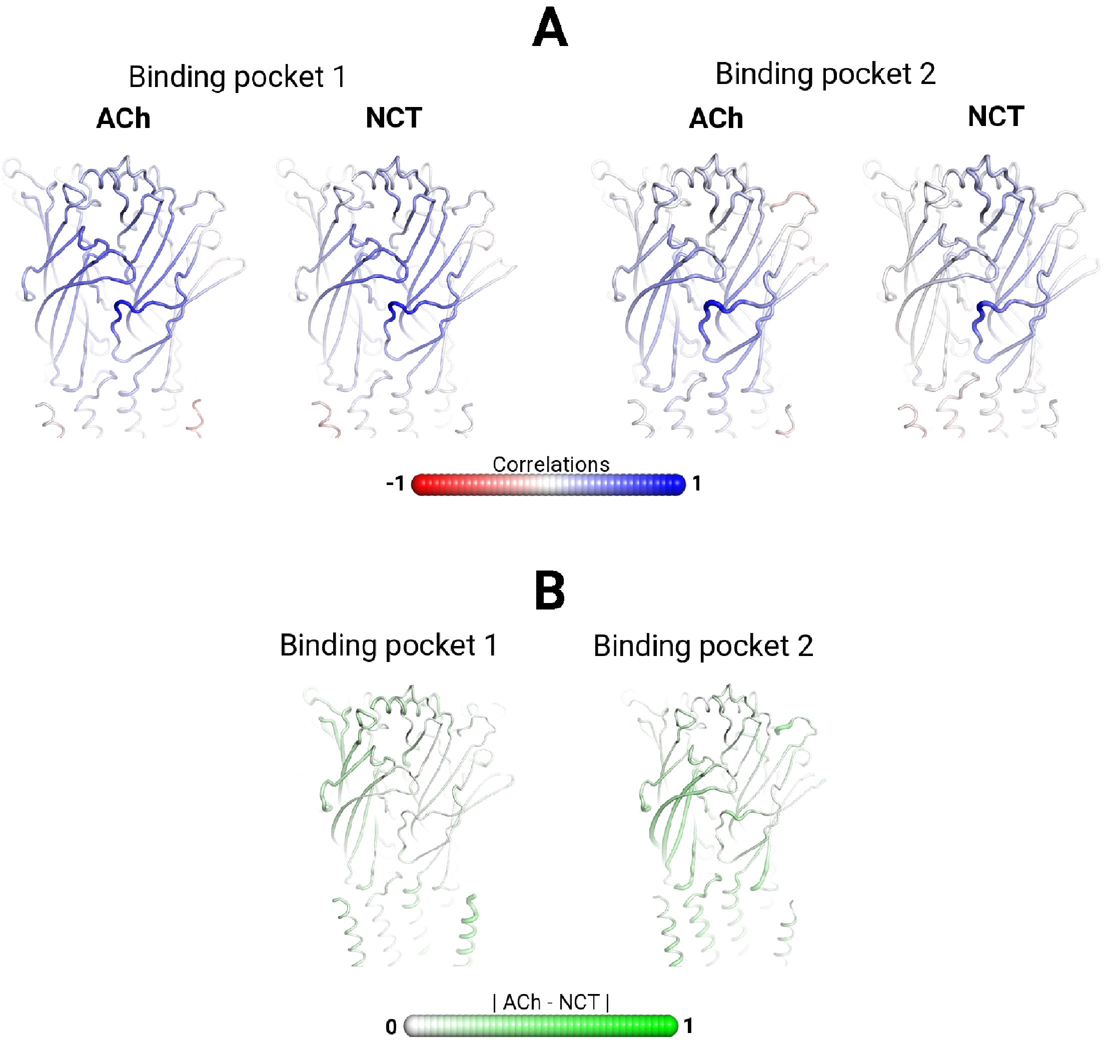

**Figure S19.** (**A**) Statistical correlation between the Cα atom of D171 (located in the top part of loop F) in the β2 subunit and all the remaining Cα atoms for the ACh- and NCT-bound systems. In this panel, the atoms that systematically move along the same/opposite direction have a correlation value of 1/-1, whereas those whose movements are uncorrelated present a correlation value of 0. (**B**) Difference between the correlations for the ACh- and NCT-bound systems.

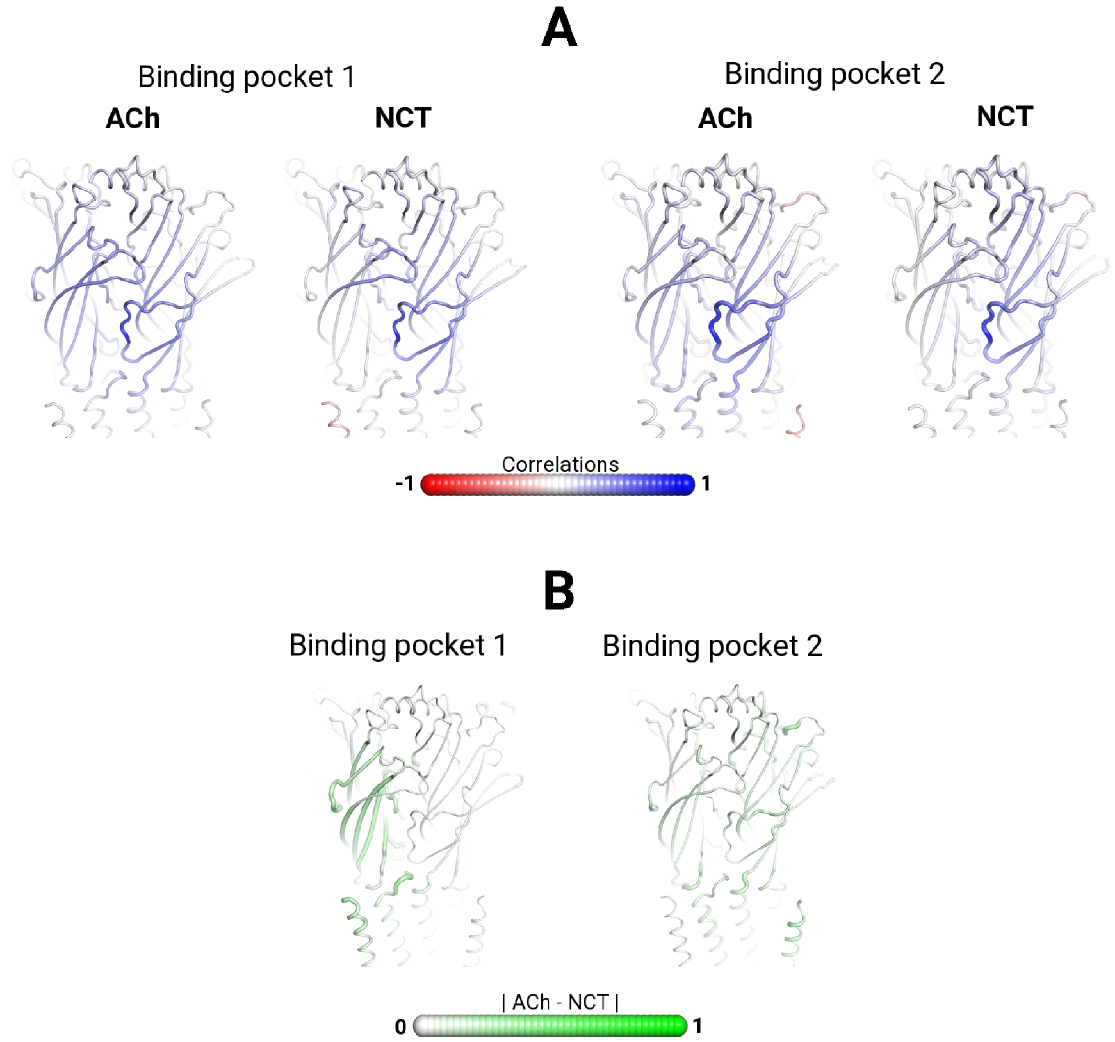

**Figure S20.** (**A**) Statistical correlation between the Cα atom of P174 (located in the lower part of loop F) in the β2 subunit and all the remaining Cα atoms for the ACh- and NCT-bound systems. In this panel, the atoms that systematically move along the same/opposite direction have a correlation value of 1/-1, whereas those whose movements are uncorrelated present a correlation value of 0. (**B**) Difference between the correlations for the ACh- and NCT-bound systems.

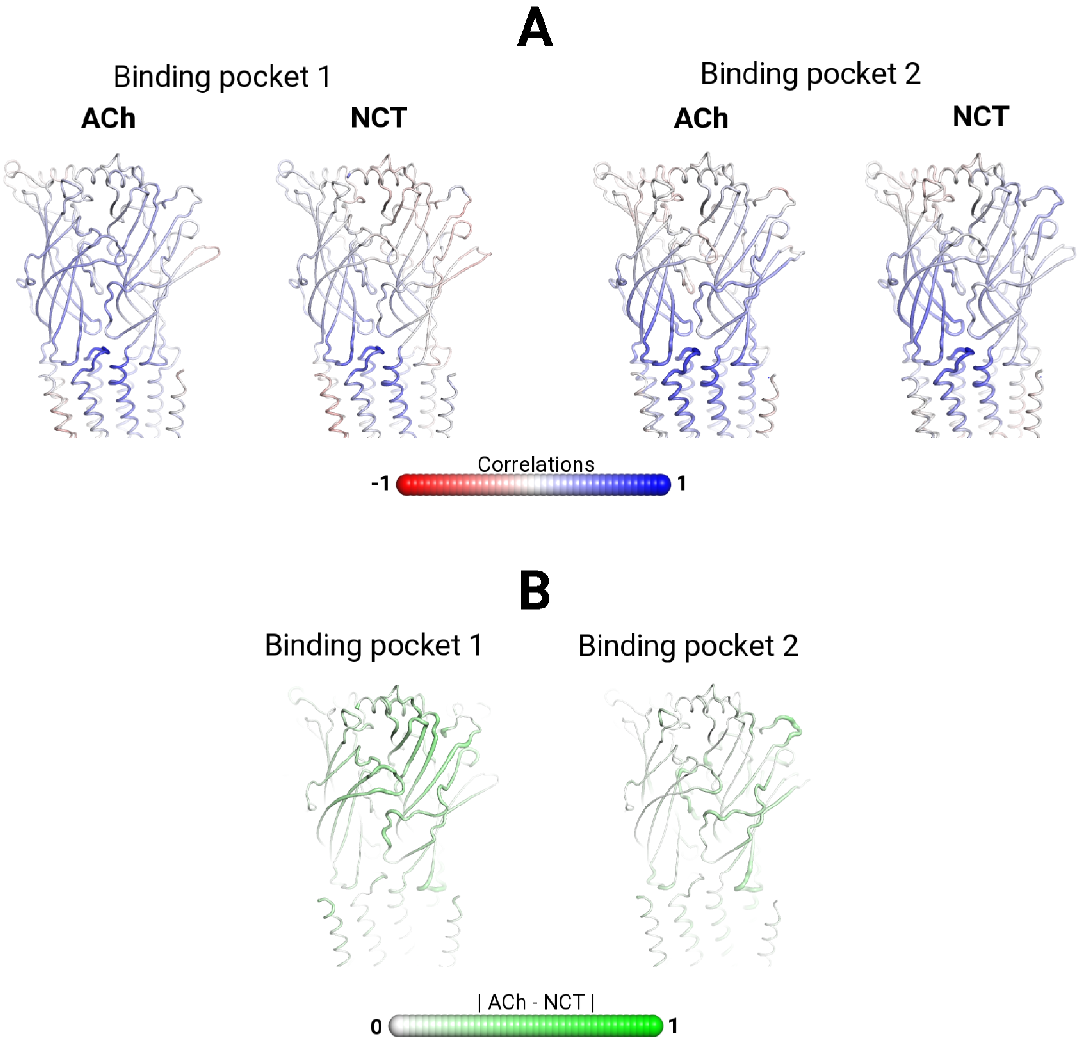

**Figure S21.** (**A**) Statistical correlation between the Cα atom of S272 (located in the M2-M3 linker) in the α4 subunit and all the remaining Cα atoms for the ACh- and NCT-bound systems. In this panel, the atoms that systematically move along the same/opposite direction have a correlation value of 1/-1, whereas those whose movements are uncorrelated present a correlation value of 0. (**B**) Difference between the correlations for the ACh- and NCT-bound systems.

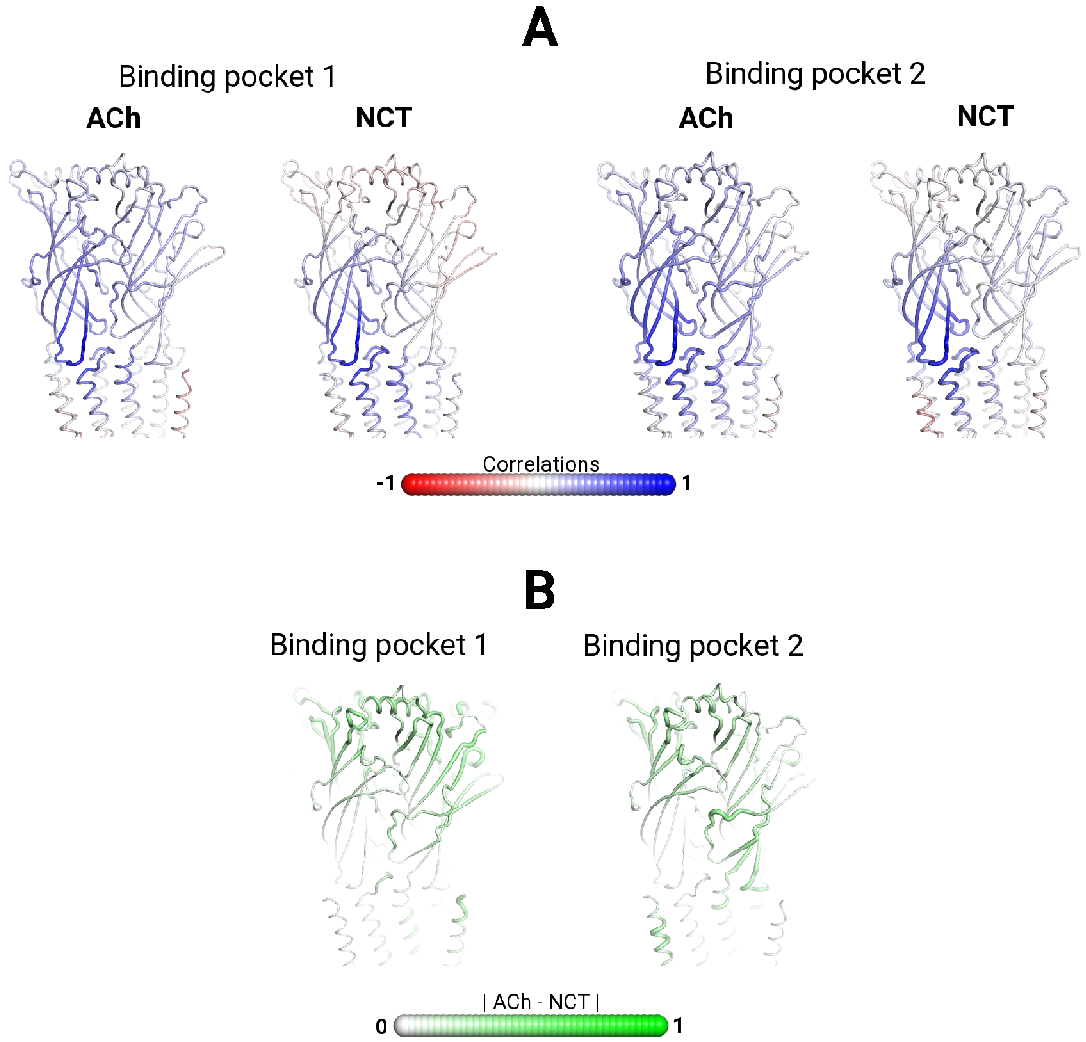

**Figure S22.** (**A**) Statistical correlation between the Cα atom of D138 (located in Cys loop) in the α4 subunit and all the remaining Cα atoms for the ACh- and NCT-bound systems. In this panel, the atoms that systematically move along the same/opposite direction have a correlation value of 1/-1, whereas those whose movements are uncorrelated present a correlation value of 0. (**B**) Difference between the correlations for the ACh- and NCT-bound systems.

#### Dynamic and structural changes in the ion channel

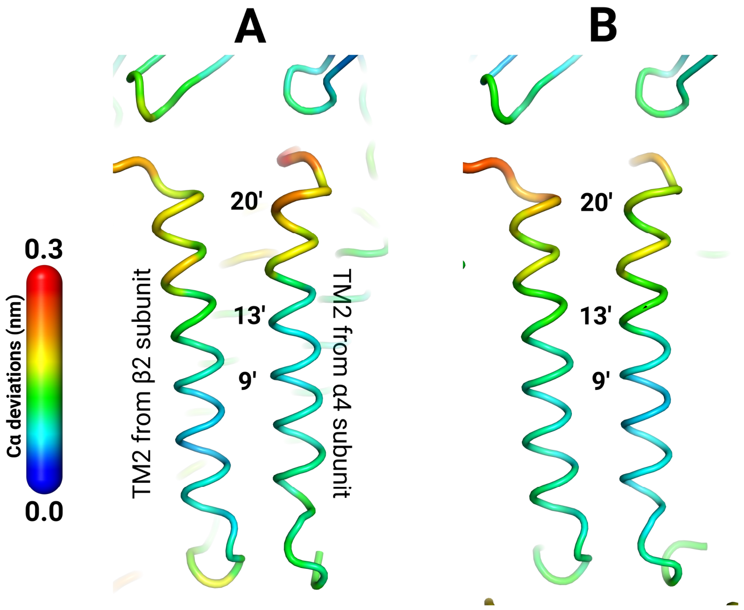

**Figure S23. Detailed view of structural differences in TM2 helices in the first (A) and second (B) α4β2 interfaces.** The TM2 helices form the inner wall of the ion channel. The hydrophobic gate is formed by the leucine and valine residues located in positions 9' and 13'. The average Cα-positional deviation between the ACh- and NCT-bound systems is mapped on the average ACh structure.

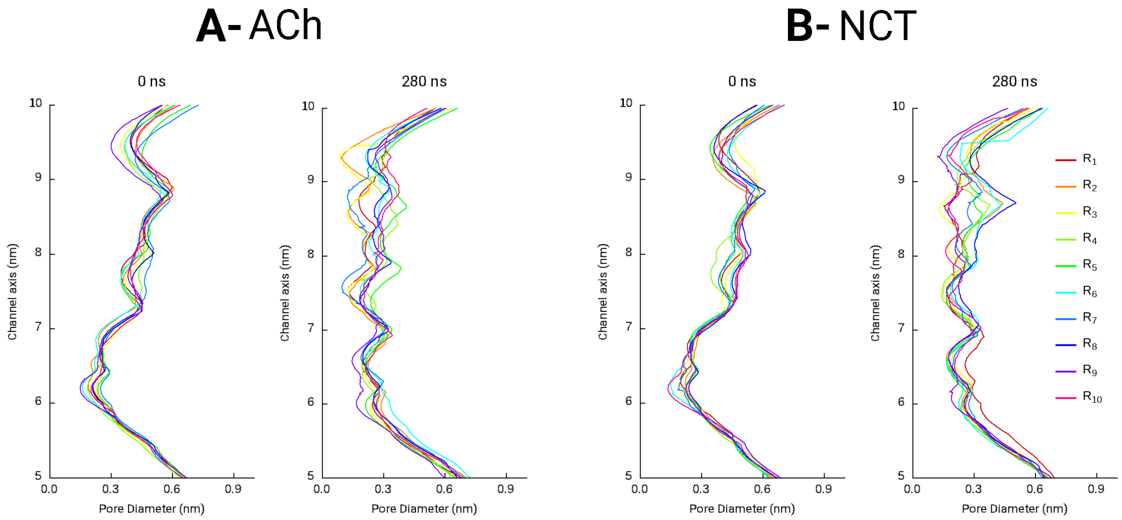

**Figure S24.** Pore diameter for the individual replicates at the beginning (t= 0 ns) and after 280 ns of simulation for the ACh- (**A**) and NCT-bound (**B**) systems. The profiles were determined with the program HOLE (*30*). For a detailed view, please zoom into the image.

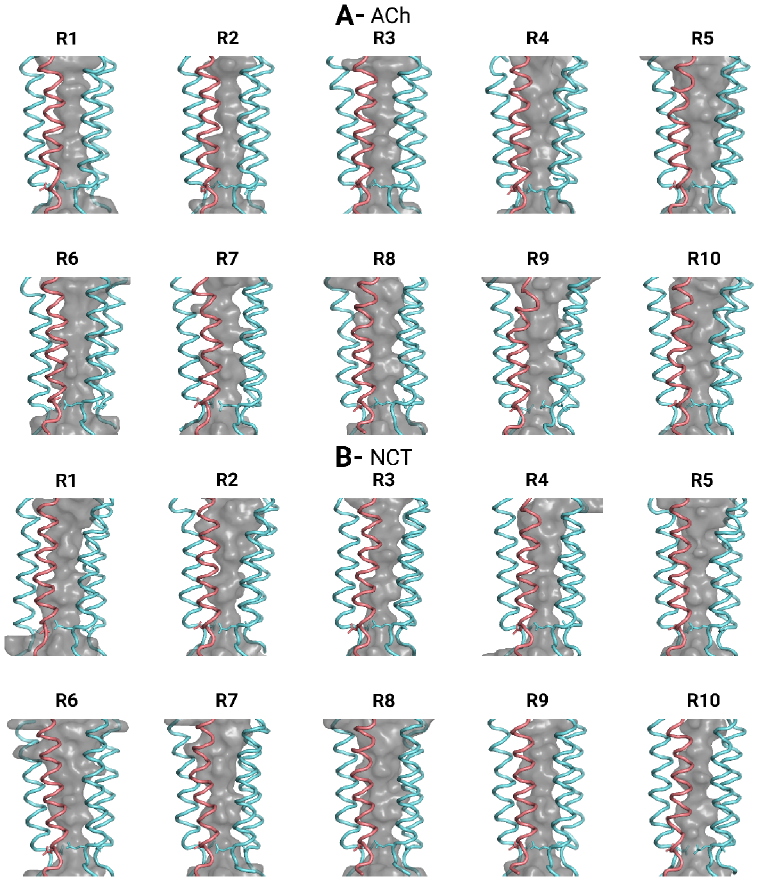

**Figure S25.** Ion permeation channel after 280 ns for all the replicates in the ACh (**A**) and NCT-bound (**B**) systems. In this image and for simplicity reasons, only the TM2 helices are depicted. The side chain of E247 (α4 subunit) and E239 (β2 subunit) are represented with sticks. The α4 and β2 subunits are coloured in pink and cyan, respectively. The grey surface corresponds to the internal surface of the transmembrane ion channel. For a detailed view, please zoom into the image.

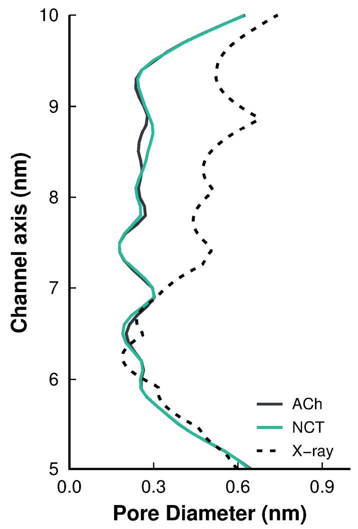

**Figure S26.** Average pore diameter after 280 ns of simulation for the ACh- (solid black line) and NCT-bound (solid green line) systems and the X-ray structure used as starting point for the simulations (*1*) (dashed line).

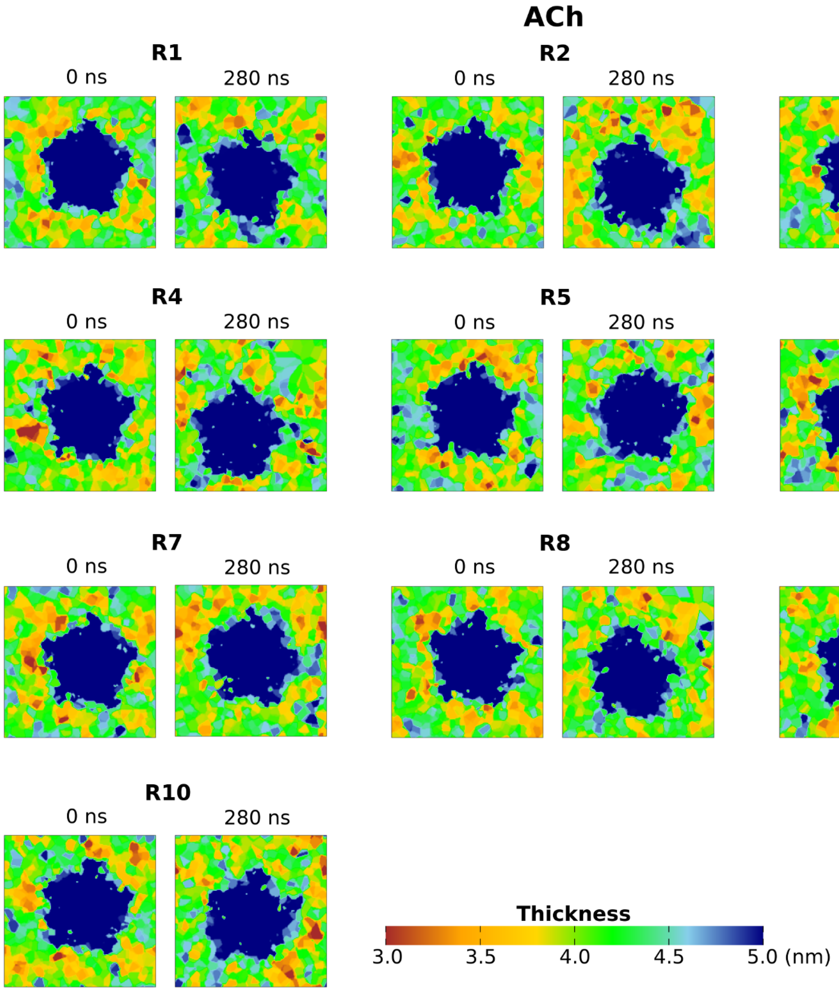

**Figure S27.** Membrane thickness at the beginning (t= 0 ns) and in the end (t= 280 ns) of the simulations for the Ach-bound systems. The thickness was calculated using the Grid-MAT tool (*31*) and projected onto the xy plane. In this image, the dark blue coloured regions indicate the location of the protein.

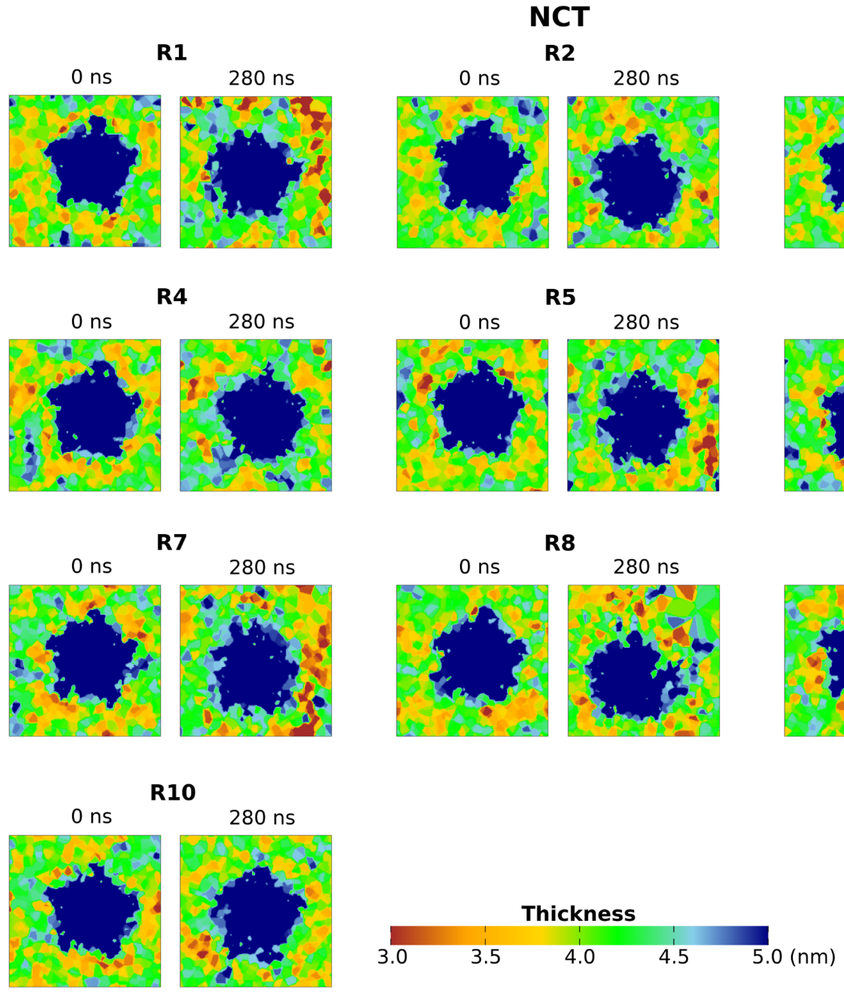

**Figure S28.** Membrane thickness at the beginning (t= 0 ns) and in the end (t= 280 ns) of the simulations for the NCT-bound systems. For details, see Figure S27.

### D-NEMD analysis

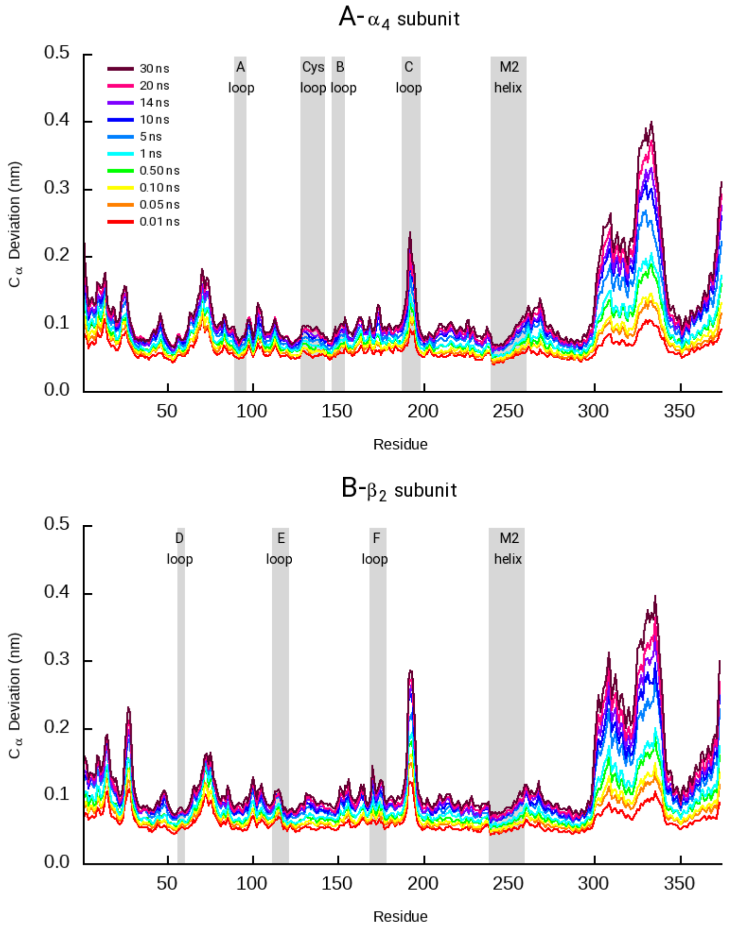

**Figure S29.** Average Cα-positional deviation for the 30 ns after ACh removal from the first binding pocket. The average deviation was calculated using the Kubo-Onsager approach (*21-24*) for the comparison of the 410 nonequilibrium free (no agonist-bound) with the correspondent equilibrium ACh-bound simulations. The positions of some essential structural motifs are highlighted in grey. For a detailed view, please zoom into the image.

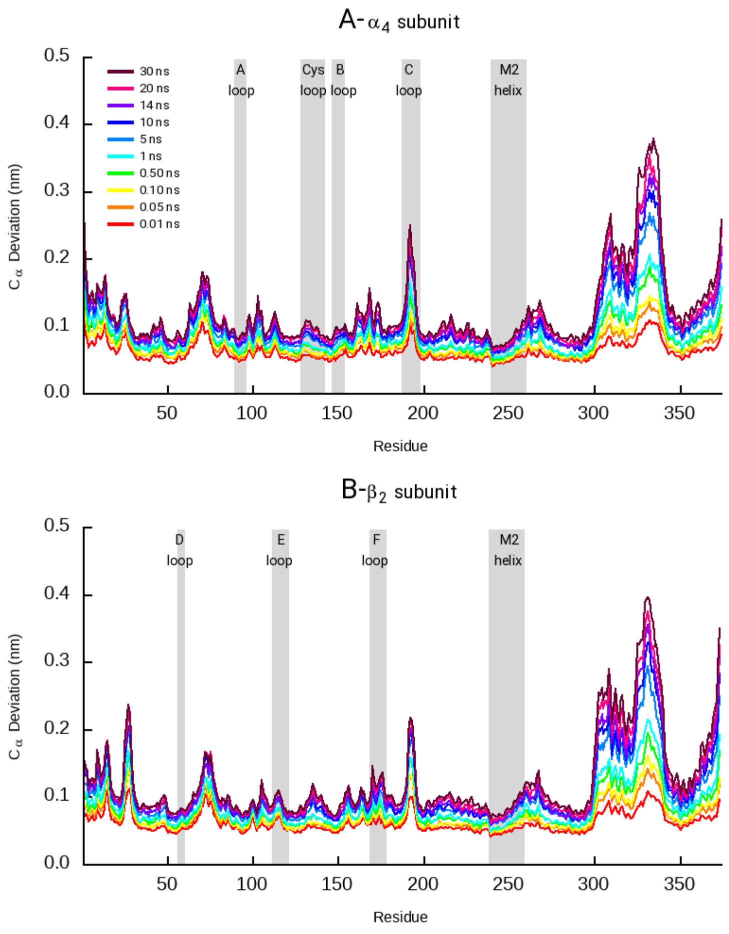

**Figure S30.** Average Cα-positional deviation for the 30 ns after ACh removal from the second binding pocket. For details, see Figure S29. For a detailed view, please zoom into the image.

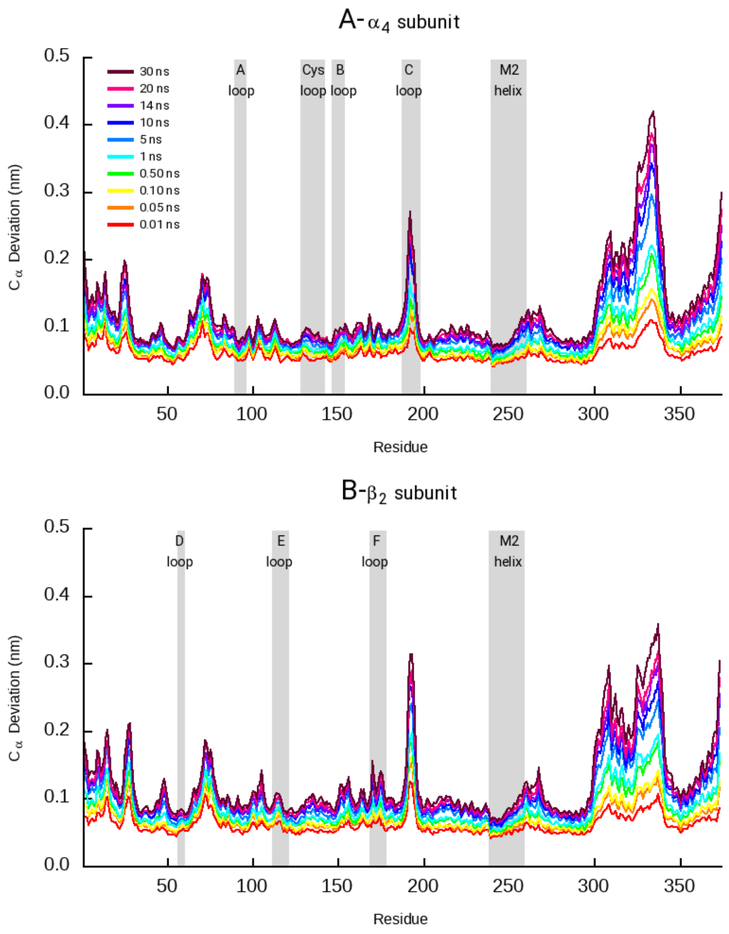

**Figure S31.** Average Cα-positional deviation for the 30 ns after NCT removal from the first binding pocket. The average deviation was calculated using the Kubo-Onsager approach (*21-24*) for the comparison of the 410 nonequilibrium free (no agonist-bound) with the correspondent equilibrium NCT-bound simulations. The positions of some essential structural motifs are highlighted in grey. For a detailed view, please zoom into the image.

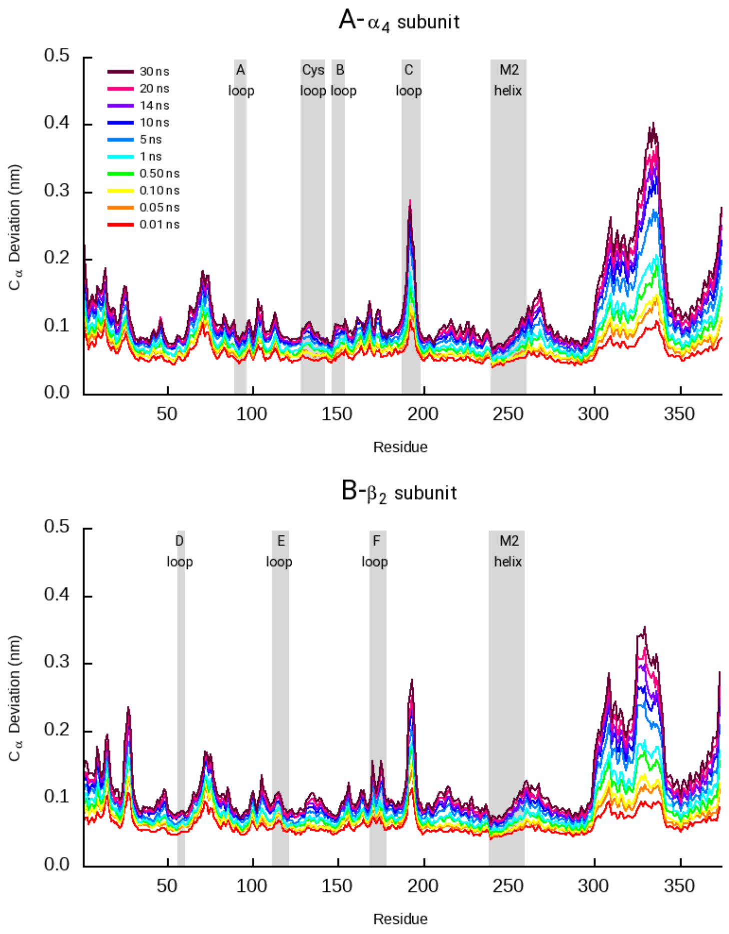

**Figure S32.** Average Cα-positional deviation for the 30 ns after NCT removal from the second binding pocket. For details, see Figure S31. For a detailed view, please zoom into the image.

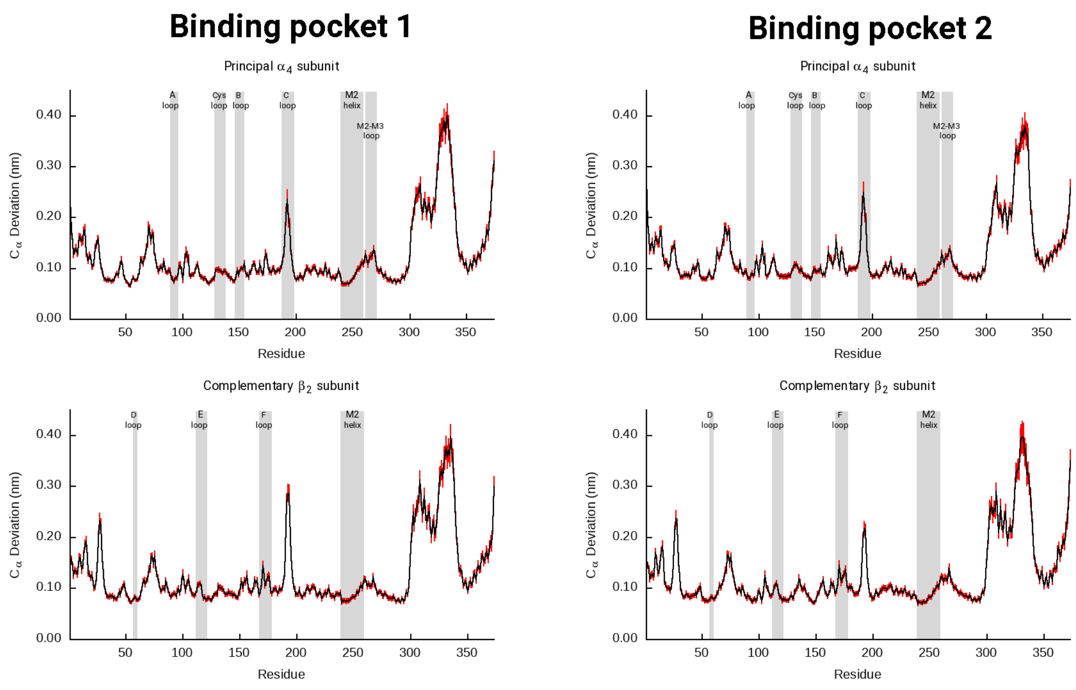

**Figure S33.** Average Cα-positional deviation 30 ns upon ACh removal from the first (right-side images) and second (left-side images) binding pockets. The vertical red lines represent the standard error of the mean. The average deviations were calculated using the Kubo-Onsager approach (*21-24*) for the comparison of the 410 nonequilibrium free (no agonist-bound) with the correspondent equilibrium ACh-bound simulations. For a detailed view, please zoom into the image.

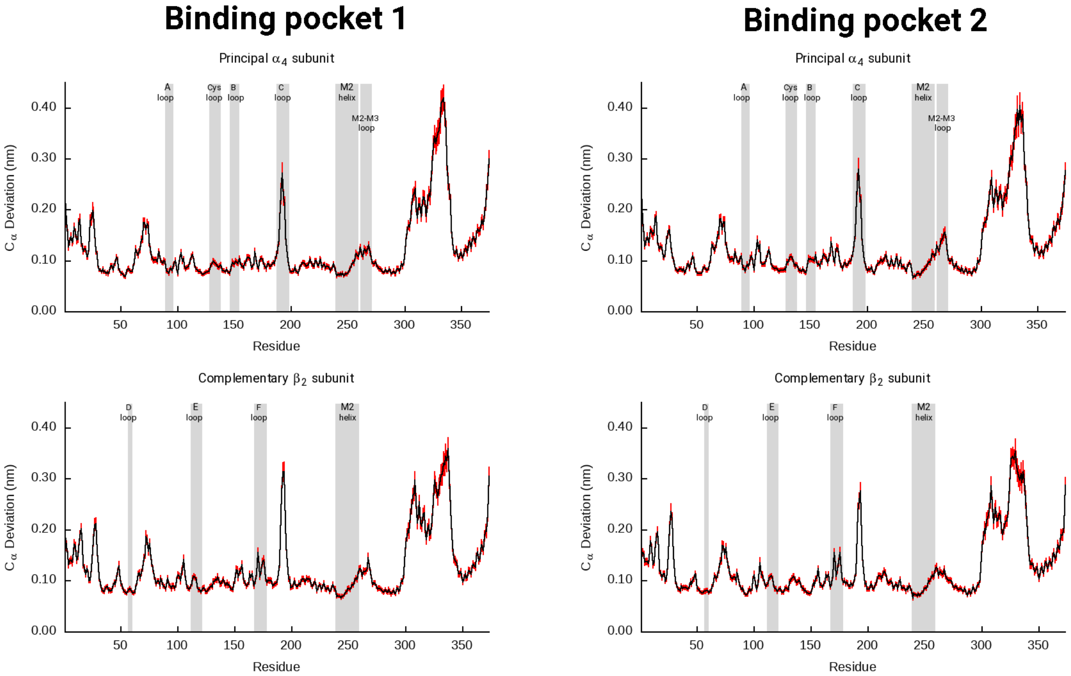

**Figure S34.** Average Cα-positional deviation 30 ns upon NCT removal from the first (right-side images) and second (left-side images) binding pockets. The vertical red lines represent the standard error of the mean. The average deviations were calculated using the Kubo-Onsager approach (*21-24*) for the comparison of the 410 nonequilibrium free (no agonist-bound) with the correspondent equilibrium NCT-bound simulations. For a detailed view, please zoom into the image.

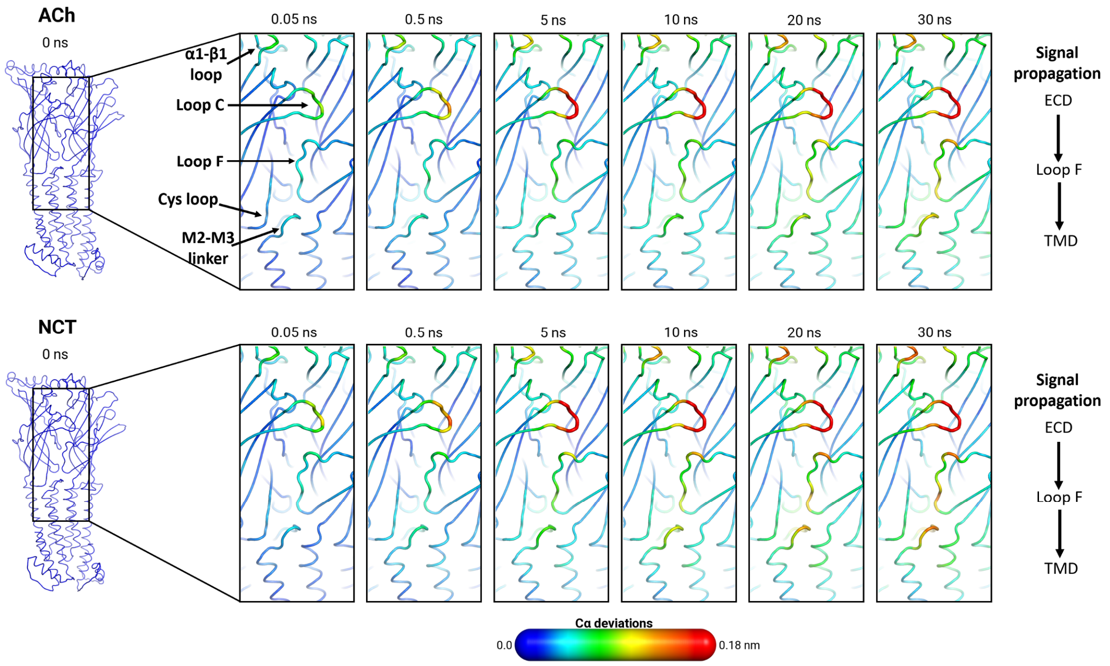

**Figure S35.** Mapping of the average Cα-positional deviation in the 30 ns following agonist removal from the second binding pocket. The $C_\alpha$ deviations between the nonequilibrium agonist-free and the equilibrium agonist-bound simulations at specific times (0, 0.05, 0.5, 5, 10, 20 and 30 ns) upon ligand removal were calculated as a function of the residue number. The final deviation values correspond to the average obtained over all 410 pairs of simulations (Figure S29 and S31) and are mapped on the average agonist-free structure using the colour scheme presented in the scale.

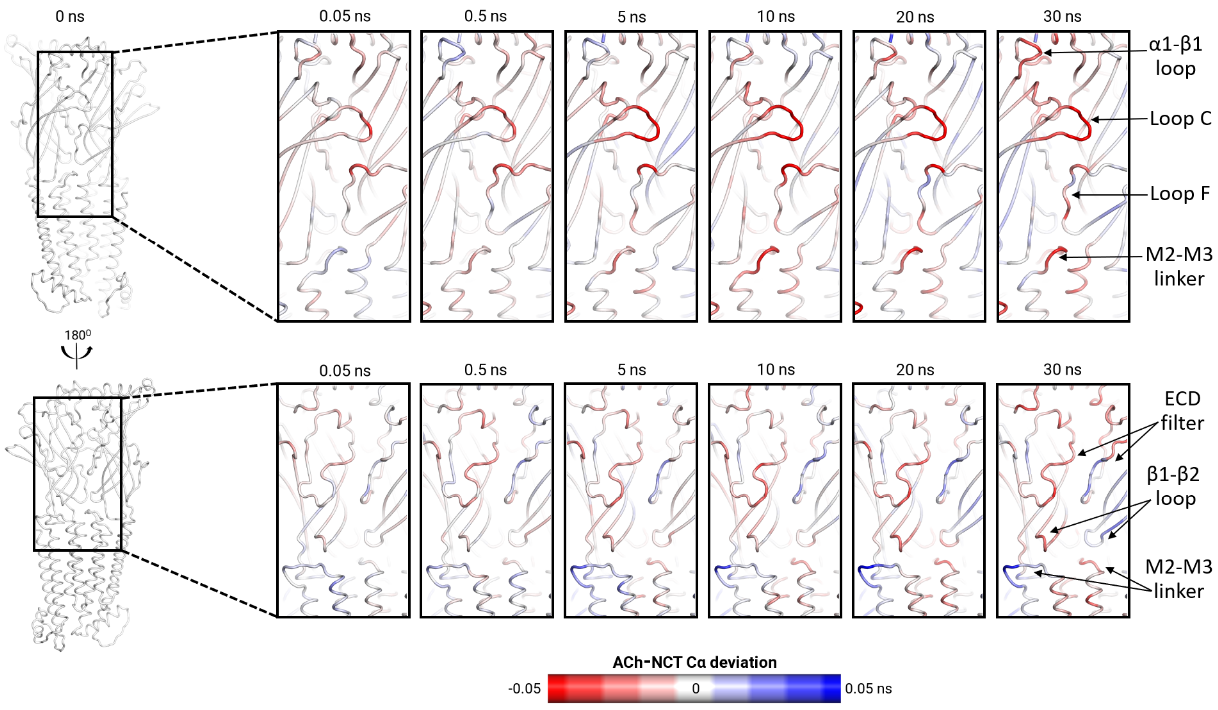

**Figure S36. Differences in response to agonist removal between the ACh- and NCT-bound systems.** The difference in the average Cα positional deviation between the ACh- and NCT-bound systems in the 30 ns following agonist removal from the second binding pocket is mapped on the average agonist-free structure.

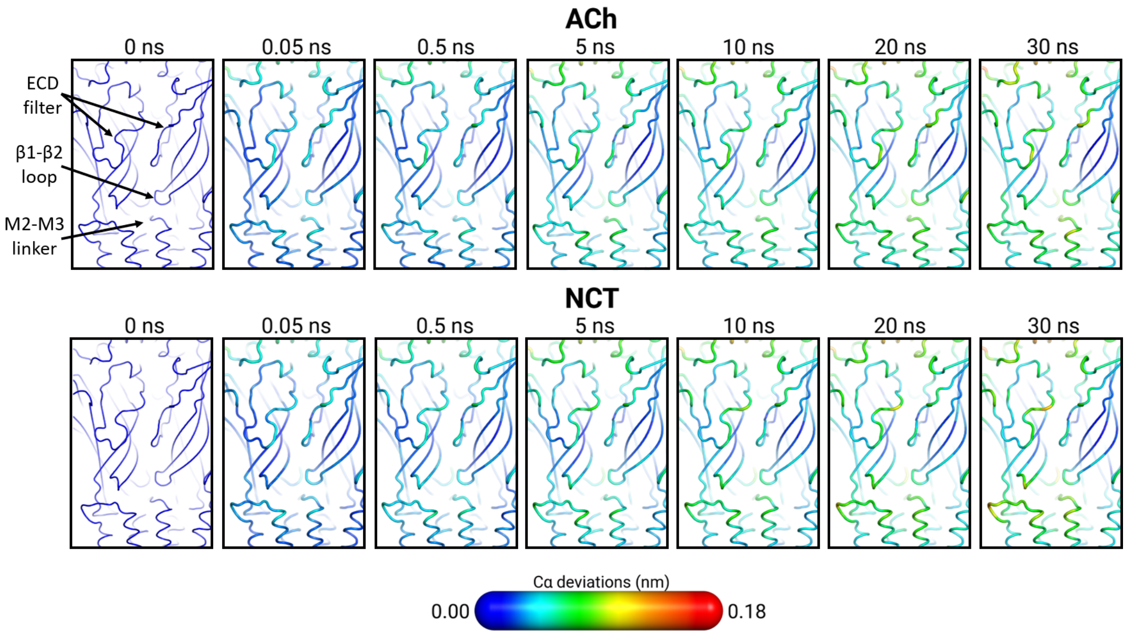

**Figure S37.** Signal propagation in the ECD selectivity filter region. Mapping of the average Cα-positional deviation in the 30 ns following agonist removal from the first binding pocket. For details, see Figure S35.

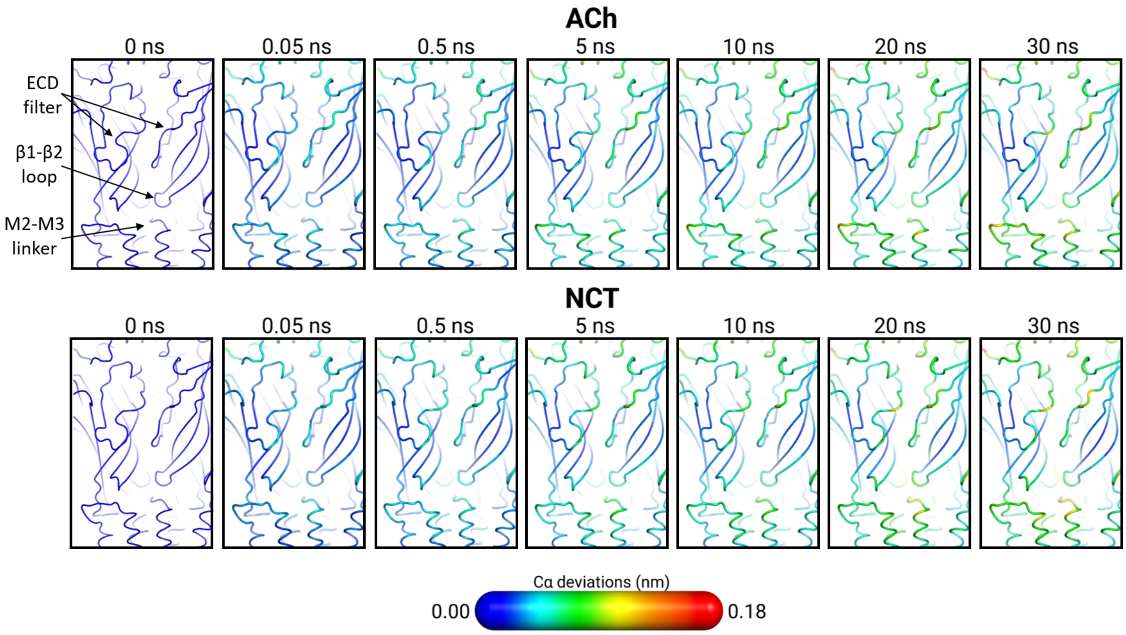

**Figure S38.** Signal propagation in the ECD selectivity filter region. Mapping of the average Cα-positional deviation in the 30 ns following agonist removal from the second binding pocket. For details, see Figure S35.

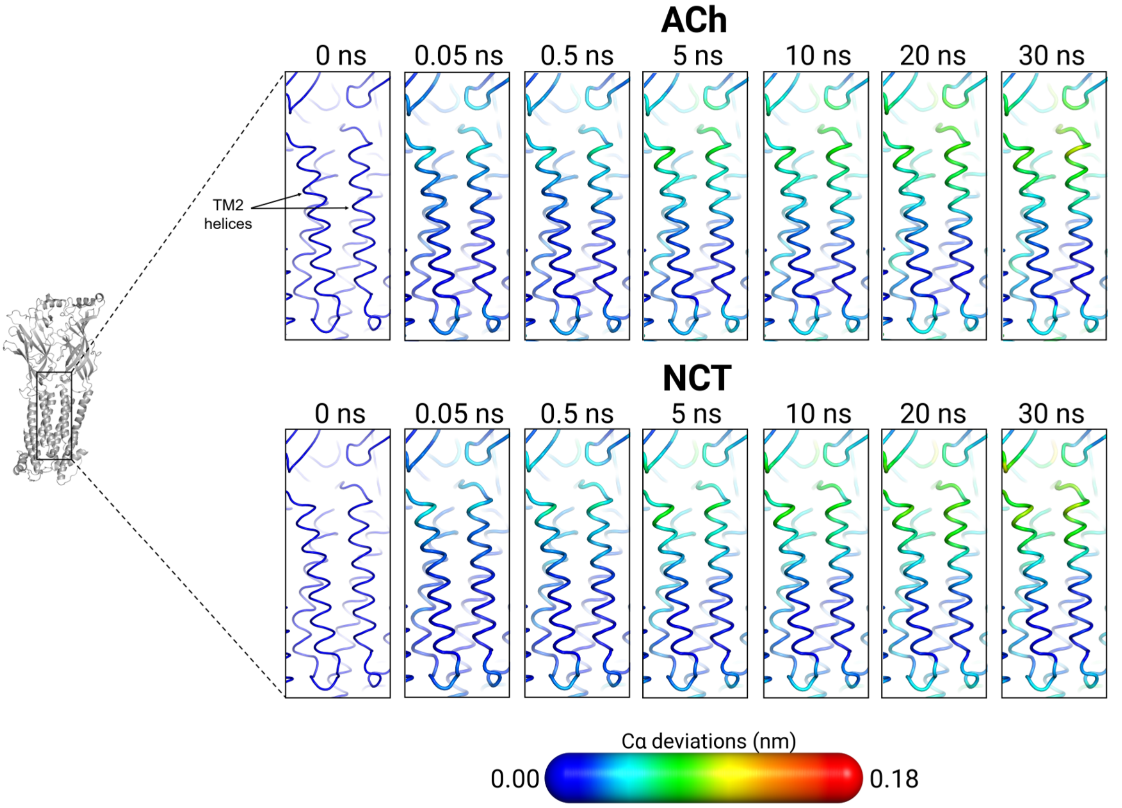

**Figure S39.** Signal propagation in the TM2 helix. Mapping of the average Cα-positional deviation in the 30 ns following agonist removal from the first binding pocket. For details, see Figure S35. Please zoom into the image for detailed visualisation.

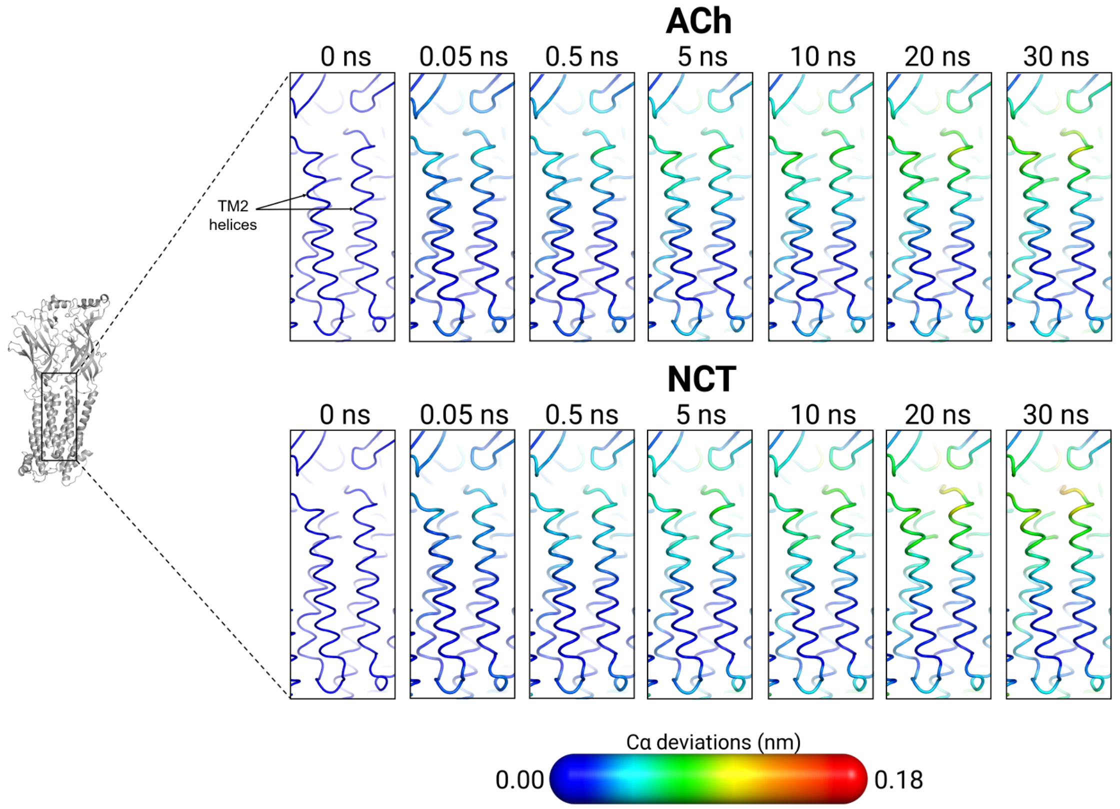

**Figure S40.** Signal propagation in the TM2 helix. Mapping of the average Cα-positional deviation in the 30 ns following agonist removal from the second binding pocket. For details, see Figure S35. Please zoom into the image for detailed visualisation.

#### Movie 1

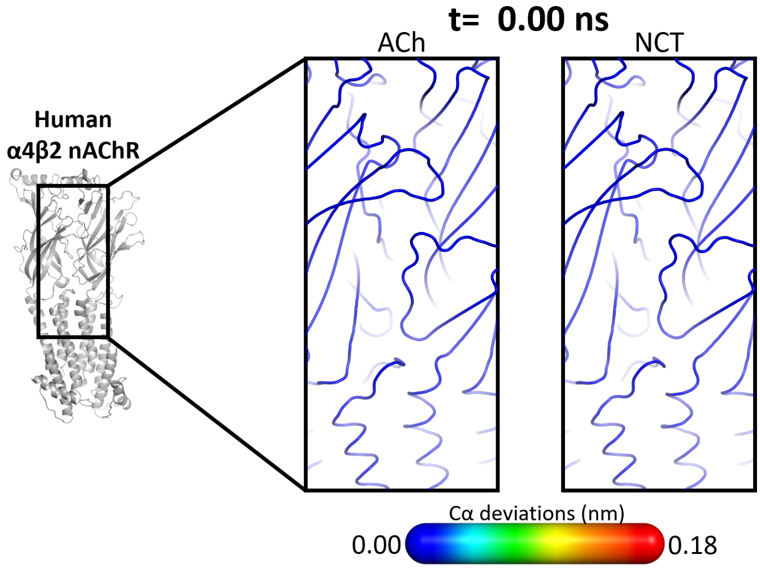

**Movie 1-** Movie depicting the time evolution of the deviations in the ECD/TMD interface over the 30 ns after agonist deletion.

#### Movie 2

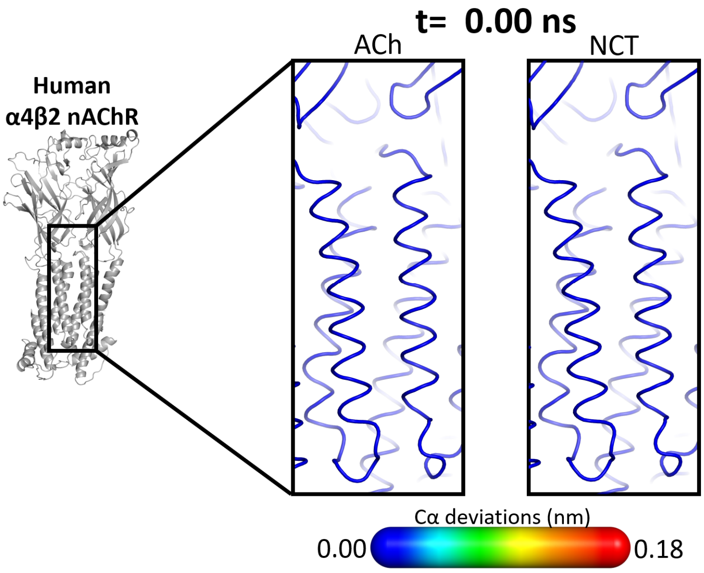

**Movie 2-** Movie depicting the time evolution of the deviations in the TM2 helices over the 30 ns after agonist deletion.
